# Cancer cells are uniquely susceptible to accumulation of MMBIR mutations

**DOI:** 10.1101/2020.07.19.209445

**Authors:** Beth Osia, Thamer Alsulaiman, Tyler Jackson, Juraj Kramara, Suely Oliveira, Anna Malkova

## Abstract

Microhomology-mediated break-induced replication (MMBIR) is a mechanism of polymerase template switching at microhomology, which can produce complex genomic rearrangements (CGRs), underlies neurological and metabolic diseases, and contributes to cancer development. Yet, the extent of MMBIR activity in genomes is poorly understood due to difficulty in directly identifying MMBIR events by whole genome sequencing (WGS). Here, by using our newly developed MMBSearch software, we directly detect MMBIR events in human genomes and report substantial differences in frequency and complexity of MMBIR events between normal and cancer cells. MMBIR events appear only as germline variants in normal human fibroblast cells but readily accumulate *de novo* across several cancer types. Detailed analysis of MMBIR mutations in lung adenocarcinomas revealed MMBIR-initiated chromosome fusions that disrupted potential tumor suppressor genes and induced CGRs. Our findings document MMBIR as a trigger for widespread genomic instability and highlight MMBIR as a potential driver of tumor evolution.

## Introduction

Massive chromosomal rearrangements are hallmarks of cancer and can promote a number of other diseases as well. The discovery of chromothripsis, which is associated with various types of cancer, demonstrated that chromosomal rearrangements can be highly complex, and that they are often induced by a single, catastrophic event (Berger et al., 2011; Kloosterman et al., 2011; Kloosterman et al., 2012; Leibowitz, Zhang, & Pellman, 2015; Stephens et al., 2011; Zhang, Leibowitz, & Pellman, 2013). This represented a significant shift in our understanding of chromosome instability, which had previously been believed to represent accumulation of individual, small genome changes over time. To explain the nature of the initial event triggering such catastrophes, two mechanisms were proposed. First, the massive shattering of a chromosome followed by random stitching together of fragments by non-homologous end-joining (NHEJ) resulting in chromothriptic chromosomes was supported by analysis of cancer genomes by Illumina sequencing (Berger et al., 2011; Kloosterman et al., 2011; Stephens et al., 2011). Alternatively, it has been proposed that chromothripsis may represent abnormal DNA synthesis that proceeds by multiple rounds of template switching to produce complex genome rearrangements (CGRs) and sometimes copy number gains (Carvalho & Lupski, 2016; Leibowitz et al., 2015; Liu et al., 2011; Zhang et al., 2015).

The latter hypothesis was originally formulated based on analysis of CGRs associated with several congenital neurological disorders, which revealed that the genetic rearrangements at CGR break sites are highly complex (Beck et al., 2015; Beck et al., 2019; Carvalho et al., 2013; Lee, Carvalho, & Lupski, 2007). Specifically, Sanger sequencing analysis of the segmental duplications thought to cause Pelizaeus–Merzbacher disease (PMD) identified complex combinations of break points, which presented evidence of template switching at microhomologies that led to copy number gains from duplications up to quadruplications (Beck et al., 2015; Lee et al., 2007). The authors proposed that this mechanism, named microhomology-mediated break-induced replication (MMBIR)(Hastings, Ira, & Lupski, 2009; Hastings, Lupski, Rosenberg, & Ira, 2009; Payen, Koszul, Dujon, & Fischer, 2008), underlies a number of neurological disorders, and represents an alternative to NHEJ in triggering chromothripsis in cancer (Liu et al., 2011). Initial Illumina sequencing of chromothriptic oncogenomes did not identify templated insertions at breakpoints indicative of MMBIR (Berger et al., 2011; Kloosterman et al., 2011; Stephens et al., 2011). However, a recent study of chromothriptic genomes selected from over 2,500 cancers, which also used more sophisticated analysis methods, revealed templated insertions at some chromothriptic junctions that could be explained by MMBIR (Cortes-Ciriano et al., 2020).

Despite this recent progress, the frequency of MMBIR and how significantly it may contribute to chromothripsis remain poorly understood. Currently, our knowledge of MMBIR is based on characterization of the junctions of CGRs or copy number variations (CNVs) (Arlt, Rajendran, Birkeland, Wilson, & Glover, 2012; Beck et al., 2015; Carvalho et al., 2013; Carvalho et al., 2015; Conrad et al., 2010; Cortes-Ciriano et al., 2020; Lee et al., 2007; Y. Wang et al., 2015; Zhang et al., 2015). This limits our understanding to only those MMBIR events that result in CGRs, and even in these cases, some MMBIR events are missed due to alignment problems. It is likely that MMBIR may occur more frequently but results in small-scale genetic lesions that are not detectable using current approaches. To advance our understanding of the contribution of MMBIR to genetic stability and genome evolution, a forward approach with the resolution to detect individual and non-catastrophic MMBIR events that do not produce CGRs is required.

Previously, we defined the genetic signature of MMBIR in our disomic yeast system, where we demonstrated that MMBIR can be initiated by the interruption of break-induced replication (BIR) (Sakofsky et al., 2015), a DSB repair pathway initiated by invasion of a broken DNA end into a homologous template followed by copying large chromosomal regions (Figure 1A), reviewed in (Kramara, Osia, & Malkova, 2018; Sakofsky & Malkova, 2017). Our work demonstrated that deletion of *Pif1* helicase, which is required for processive BIR replication, promoted template switching at sites of microhomology leading to MMBIR. From these studies, we established a signature of MMBIR defined by a short insertion copied from a nearby template (within 100 bp from the position of insertion) and flanked by microhomology (Sakofsky et al., 2015).

**Figure 1.**
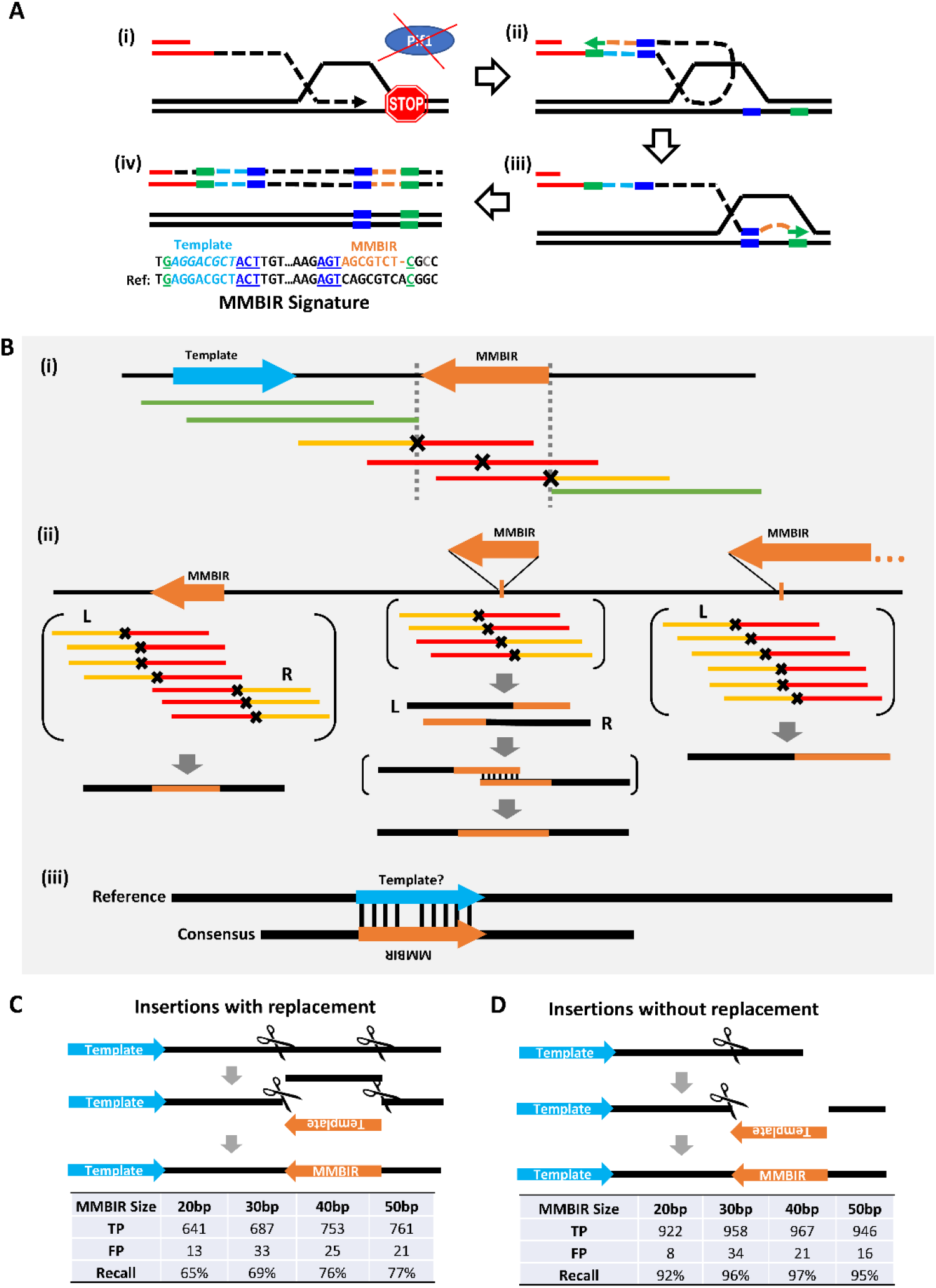
MMBSearch is a novel tool to detect MMBIR events in NGS reads. **(A)** Schematic of the MMBIR mechanism from ^22^ where **(i)** BIR collapses; **(ii)** The 3’ end of the nascent strand dissociates from its template and anneals at microhomology (dark blue rectangle) and copies (orange dotted line) the new template (light blue dotted line). **(iii)** The newly synthesized nascent strand dissociates from the template and anneals at microhomology (green rectangle) to another template where BIR continues. **(iv)** The outcome (signature) of MMBIR (top): insertion (orange) located in proximity and inverted orientation to its complementary template (light blue) and flanked by microhomologies at the junctions. Bottom: the original sequence (reference). **(B)** The MMBSearch tool identifies MMBIR events based on the signature described in A: **(i)** Reads containing MMBIR insertions (red), which do not align to a reference genome are collected and re-aligned as half-reads. **(ii)** Reads where the first half (yellow) aligns while the second half (red) does not are collected and clustered by position of the aligned halves. The aligned read halves serve as anchors, and whole reads anchored on the same side (Right – R or Left - L) are used to create consensus sequences. If R and L consensuses overlap by position and are discordant (middle example), they are aligned to each other to create the full accurate consensus, which is compared back to the reference genome to identify the full MMBIR insertion sequence. **(iii)** The reverse-complement of the MMBIR insertion is aligned to the reference genome within 100 bp from the insertion to identify a possible template. **(C) (top)** Schematic for creation of synthetic insertions by replacing sequence with the reverse complement of a template from 80bp upstream (5’ of the insertion). **(bottom)** Number of true positives (TP), number of false positives (FP) and recall (TP/total) called by MMBSearch from processing of synthetic reads generated from Chromosome 17 with 993 synthetic insertions sized 20-50bp. **(D) (top)** Schematic for creation of synthetic insertions by appending the reverse complement of a template from 80bp upstream **(bottom)** Analysis for the reads in **top** analyzed similarly to **C.**

Here, we describe the utilization of this MMBIR signature to create a new software tool called MMBSearch, which uses a highly sensitive mapping strategy to identify MMBIR insertions from whole-genome sequencing (WGS) reads irrespectively of whether they are associated with CNVs or CGRs. Using our MMBSearch software, we report the first genome-wide characterization of MMBIR in normal and cancer human cells. We observed accumulation of MMBIR events in several human cancers. By contrast, MMBIR events in normal human skin fibroblasts were present only as germline events and did not accumulate in somatic cells. Sequencing of non-small-cell lung cancer tumors identified MMBIR events that initiated a chromothripsis-like pattern of CGRs. Our data demonstrate that MMBSearch is a powerful tool that can be used effectively in multiple cell contexts to analyze the frequency of MMBIR and the associated genetic outcomes.

## Results

### MMBSearch identifies insertions with nearby templates in NGS datasets

To identify all MMBIR events genome wide, we developed a computational search tool, MMBSearch, that uses an MMBIR signature determined from our studies in yeast (Sakofsky et al., 2015) (Figure 1A). Specifically, MMBSearch identifies small templated insertions (10 bp at minimum and the full length of the reads at maximum) that represent inverted copies of their templates located within 100 bp from the insertion. Because standard genomics pipelines often exclude insertions comparable in length to the read size, due either to soft-clipping or ignoring reads that do not adequately match the reference, we engineered MMBSearch to call only reads that were excluded by initial alignment, and to use a split-read alignment to cluster candidate MMBIR reads by their reference-matching half-read anchors (Figure 1Bi) (see Methods). The resulting clusters are used to create an accurate consensus from which the exact boundaries of the insertion can be determined and used to search for nearby inverse-complementary sequence representing the template (Figure 1Bii & iii).

We tested the sensitivity of MMBSearch using a set of artificially created human genomes that each contained 993 MMBIR insertions of varying lengths (from 20 to 50 bp) on human chromosome 17 (see Methods). Insertions that both replaced or did not replace the sequence at the insertion site were included (Figure 1C, D). MMBSearch analysis of artificial Illumina sequencing reads generated from these test genomes demonstrated recall (sensitivity) of 92-97% for insertions without replacement (Figure 1D) and recall of 65-77% for insertions with replacement (Figure 1C). False-positive calls were low in both cases (1-3%) (Figure 1C, D) (see Supplementary text for details). Thus, MMBSearch accurately calls MMBIR events matching the MMBIR signature from (Sakofsky et al., 2015) in human genome datasets.

### MMBIR insertions do not accumulate with age in human fibroblasts

To determine whether we could identify MMBIR events in normal somatic cells, we obtained sequencing reads originally prepared from clonal fibroblast lineages by (Saini et al., 2016). In this study, the authors assessed the frequency of mutations accumulated in an individual’s lifetime by sequencing clonal lineages derived from several locations on the forearms and hips of two individuals. The authors demonstrated that, in these samples, both base substitutions and chromosomal rearrangements accumulated at frequencies that were as high as those associated with some cancers, prompting the hypothesis that MMBIR mutations may be common and/or accumulate in these fibroblast clones as well.

To test our hypothesis, we analyzed sequencing reads from all 10 clonal fibroblast cell lineages and their matched blood samples (6 clones from individual D1 in the study and 4 clones from individual D2). Because the fibroblast lineages were clonal, we required a minimum of 10 reads per cluster to exclude any MMBIR events that may have arisen during clone expansion (see Methods for details). We set parameters for MMBSearch to call insertions with a minimum length of 10 bp and an identity threshold of 80% between the insertion and its template. MMBIR events called at the same reference positions in both fibroblasts and blood were considered germline variants (present from the zygotic stage), while those that were found only in fibroblast clones were considered fibroblast specific. Strikingly, we observed that the majority (74-93%) of all MMBSearch calls among all 10 fibroblast clones were germline variants (Table 1, Supplementary Figure 1; Supplementary Data S1, S2). The remaining 7-26% of MMBIR calls we identified as fibroblast-specific events; however, all but one call was either present among multiple fibroblast clones, represented microsatellite alterations, or were determined to be false-positive outcomes (Table 1; Supplementary Data S1, S2). The sole remaining fibroblast-specific variant was a 354-kb deletion on Chr3 in the first left forearm sample of Individual D1 (D1-L-F1) previously described in (Saini et al., 2016) (Supplementary Data S3). This deletion occurred at a short 5 bp quasi-palindrome (See Figure 2A - DQP), with just 1 bp of microhomology at the junctions. However, this event appears to differ from similar germline events (Supplementary Figure 3C; Supplementary Data S4-S7) in both size of the deletion and amount of microhomology. Therefore, it is unlikely that this event represented MMBIR. Taken together, we found no evidence of MMBIR accumulation in fibroblasts with age, which represents an intriguing difference between MMBIR mutational events and other mutations that accumulated at high frequency in aging fibroblasts (Saini et al., 2016).

**Table 1:** Summary of MMBSearch calls from 10 clonal fibroblast lineages of 2 individuals.

| Fibroblast clone ID * | All MMBSearch calls † |  | All Fibroblast-specific MMBSearch calls ‡ |  |  |  |
| --- | --- | --- | --- | --- | --- | --- |
|  | Germline calls | Fibroblast-specific calls | Alignment and adapter errors | Microsatellite alterations | Multi-clone MMBIRs | Clone-specific MMBIRs |
| D1-L-H | 1094 (82%) | 244 (18%) | 35 (14%) | 186 (76%) | 23 (9%) | 0 (0%) |
| D1-R-H1 | 1057 (87%) | 158 (13%) | 29 (18%) | 117 (74%) | 12 (8%) | 0 (0%) |
| D1-R-H2 | 1048 (84%) | 203 (16%) | 31 (15%) | 151 (74%) | 21 (10%) | 0 (0%) |
| D1-L-F1 | 1105 (78%) | 312 (22%) | 55 (18%) | 231 (74%) | 25 (8%) | 1 (0.3%) |
| D1-L-F2 | 1066 (84%) | 206 (16%) | 18 (9%) | 164 (80%) | 24 (12%) | 0 (0%) |
| D1-R-F | 998 (88%) | 130 (12%) | 25 (19%) | 92 (71%) | 13 (10%) | 0 (0%) |
| D2-L-H | 1058 (92%) | 90 (8%) | 22 (24%) | 63 (70%) | 5 (6%) | 0 (0%) |
| D2-R-H | 984 (93%) | 78 (7%) | 17 (22%) | 56 (72%) | 5 (6%) | 0 (0%) |
| D2-L-F | 1089 (85%) | 195 (15%) | 50 (26%) | 136 (70%) | 9 (5%) | 0 (0%) |
| D2-R-F | 1177 (84%) | 229 (16%) | 63 (28%) | 162 (71%) | 4 (2%) | 0 (0%) |

\* Fibroblasts were derived from individual 1 (D1) and individual 2 (D2) from the study of <sup>25</sup>. Clone sites: left hip (D1-L-H, D2-L-H), Right hip (D1-R-H1, D1-R-H2, D2-R-H), left forearm (D1-L-F1, D1-L-F2, D2-L-F) and right forearm (D1-R-F, D2-R-F).
† MMBSearch calls from clusters with $\geq 10$ reads, insertions that are $\geq 10$ bp in length, and $\geq 80\%$ identity between template and insertion.
‡ Percentages are out of total fibroblast-specific calls.

**Figure 2.**
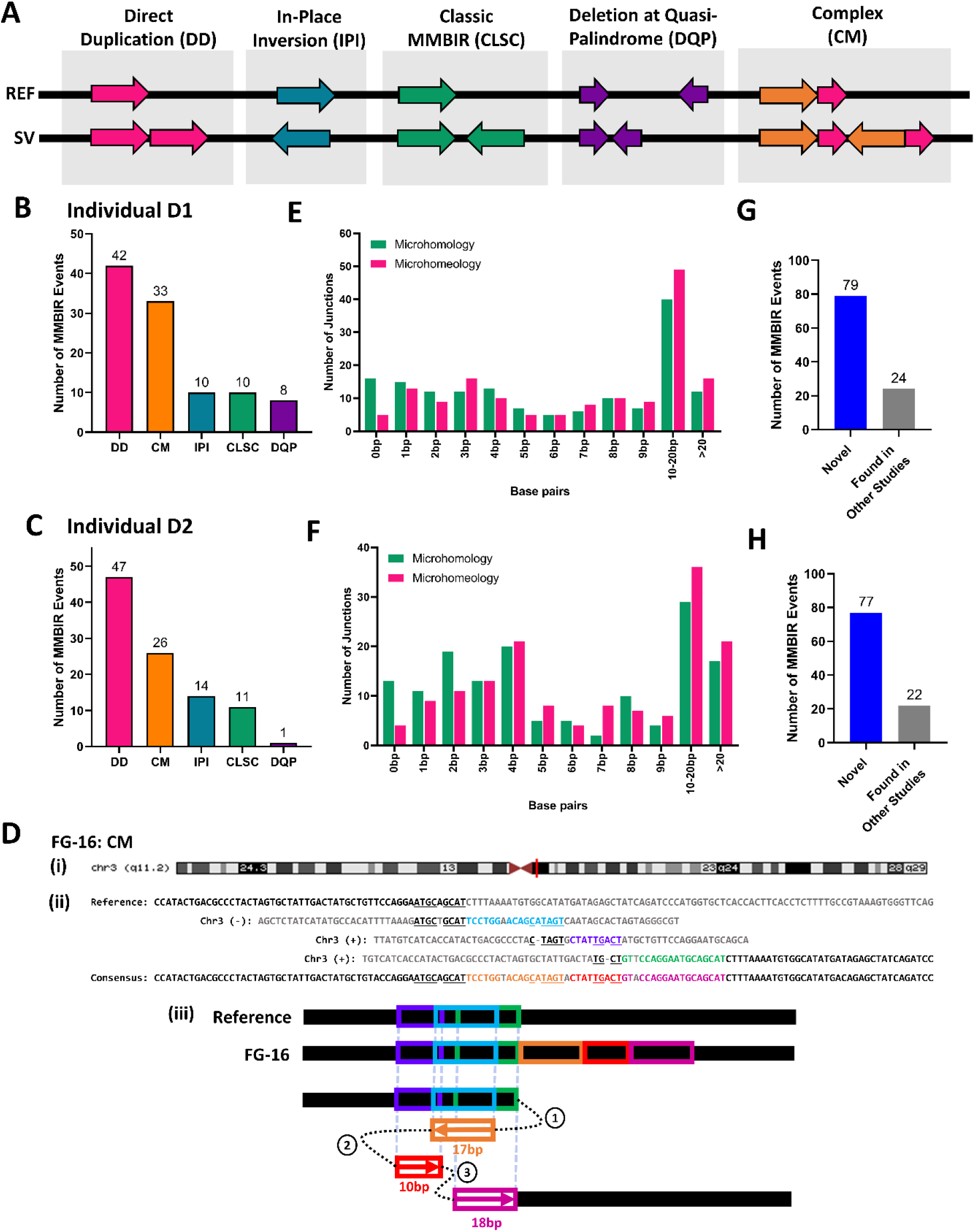
MMBIR underlies formation of germline variants found in human fibroblasts. **(A)** Schematic of different classes of MMBIR events. REF=reference; SV= MMBIR event; **(B)** Distribution of different MMBIR classes among germline events (parameters for search: ≥ 25bp in length) identified in individual D1 (n=103) ^25^. **(C)** Similar to **B**, but for individual D2 (n=99) ^25^. **(D) (i)** Complex MMBIR (CM) event FG-16 found on q11.2 band of Chromosome 3 (red vertical line indicates position; maroon triangles indicate centromere location). **(ii)** Alignment of event FG-16 consensus to all refence templates. Insertion consists of 3 parts (orange, red, and pink text) copied from all corresponding templates (light blue, purple, and green respectively). Microhomology used in each template switching event is underlined. **(iii)** Schematic illustrating template switching events that lead to the formation of FG-16. Circled numbers indicate the order of template switches. Boxed arrows indicate direction of synthesis, numbers under boxes indicate length of synthesis, and colors correspond to those in (ii). **(E)** Distribution of microhomology analyzed for all resolvable junctions of MMBIR events from D1 shown in **B**(n=155). Microhomology = identical bases; microhomeology = identical, but with 1bp mismatch or gap allowed. **(F)** Similar to **E**, but for D2 shown in **C**(n=148). **(G)** Number of novel MMBIR events from D1 (from **B**) and those already exiting in NCBI’s Nucleotide collection (nr/nt). **(H)** Similar to **G**, but for events from D2 (from **C**).

### MMBIR underlies germline insertions in normal skin fibroblasts

We next analyzed the germline variants that were the majority of fibroblast MMBSearch calls identified in the dataset from (Saini et al., 2016) (Table 1). When we set the threshold to signatures of 10 bp or longer, the number of calls was very high in both individuals (between 998 and 1105 calls in individual D1 and between 984-1177 calls for individual D2 (Table 1)), and the majority of called events represented microsatellite alterations or other noise. This level of noise was similar to what we observed among the fibroblast-specific calls using these parameters (Supplementary Data S4). To reduce the noise from shorter MMBSearch calls, we narrowed our focus to calls that were at least 25 bp (See Methods). After filtering out microsatellites and erroneous calls, we found 103 MMBsearch calls from Individual D1 and 99 calls from individual D2 that were further divided into several classes (Figure 2A; Supplementary Data S4, S6). Ten (D1) and 11 (D2) calls were classical MMBIR events, like those described in (Sakofsky et al., 2015) (Figure 2A-C (CLSC), and Supplementary Figure 2A, B). Further, 10 (D1) and 14 (D2) insertions were categorized as “in-place inversions” (Figure 2A-C (IPI) and Supplementary Figure 3A), which could either result from MMBIR-like template switching producing an insertion that completely replaces the inverted template, or from a cut-and-paste type non-homologous end joining (NHEJ) mechanism. The finding of microhomologies at the borders of all these in-place inversions supports that these resulted from an MMBIR-like mechanism (Supplementary Figure 4B (IPI); Supplementary Figure 5B (IPI)). Two additional classes of MMBIR events occurred in the regions of quasi-palindromes. The first were insertions that matched an inverted template within a quasi-palindrome that pre-existed in the reference with a few mismatches, and also matched a second direct orientation template without mismatches (thereby making it the more likely template), which we term, “direct duplications” (Figure 2A (DD); Supplementary Figure 3B). This was the most abundant class, with 42 and 47 direct duplications identified in subjects D1 and D2, respectively (Figure 2B, C). The second class were deletions that resulted from template switching inside the quasi-palindrome, which brought together two pieces of the palindrome that were not previously juxtaposed (Figure 2A (DQP); Supplementary Figure 3C). These events were much less common, with only 8 and 1 event identified in subjects D1 and D2, respectively (Figure 2B, C). Finally, we also observed cases that we termed “complex”, because they likely involved several template switching events (Figure 2A (CM) and Supplementary Figure 6). For example, one event (Figure 2D) first copied from an upstream template in the reverse direction, switched to a second template and copied in the forward direction, and then switched to a third template, copying again in the forward direction. This resulted in an insertion originating from 3 distinct templates. We identified 33 complex events in subject D1 and 26 complex events in subject D2 (Figure 2B, C).

We next explored the length of microhomologies mediating the MMBIR events uncovered by MMBSearch. For all manually verified events, we identified the junctions with resolvable templates and calculated the amount of microhomology present on their borders. We observed that 138 of 155 resolvable junctions from MMBIR events in D1 and 135 of 148 resolvable junctions from MMBIR events in D2 contained at least 1 base pair of microhomology (Figure 2E, F). In general, all MMBIR type classes contained junctions with microhomology, though microhomology length varied between types (See Supplementary Data S4-S7, and Supplementary Figures 4 & 5 for microhomology distributions by class). In addition, allowing for a single mismatch or gap to interrupt the complementary bases at junctions revealed additional bases that could have been used as microhomology during the formation of these events. We refer to this interrupted microhomology as “microhomeology,” similarly to the phenomenon described in (Anand et al., 2014; Beck et al., 2019; Carvalho & Lupski, 2016; Liu et al., 2017). Because we cannot determine whether these mismatches were present when the event formed, it is impossible to discern whether the event was initiated by annealing at a microhomeologous site or at a site of longer microhomology. In either case, the amount of microhomology that was used to promote the formation of these events could be longer than what is present at the junctions. Together, the presence of microhomologies at the resolvable junctions of the majority of germline events for both individuals indicates that they are most likely the result of template switching driven by microhomology and, therefore, we classified all outcomes in these categories (those shown in Figure 2A) as MMBIR events. Because all MMBIR events were present in both the blood and in fibroblast clones, we next asked whether they were more likely embryonic or represented common variations of evolutionary origin. However, without the ability to conduct similar analyses on sequencing from the individuals’ parents to differentiate between the two, we used BLAST to instead check whether these insertions matched other known non-reference human genome sequences. We used BLASTn to query NCBI’s nucleotide collection (nr/nt), which includes previously identified non-reference human genomic sequences, to determine whether our MMBIR events also exist in the genomes of other individuals. We found that only 24 of the 103 events from D1 and 22 of the 99 events from D2 were also found in other studies (Figure 2G, H) and therefore are not likely to be of embryonic origin, as they exist across populations of humans. The majority of the germline events that we identified in both individuals were not found in previously published studies and therefore represent undescribed events that either arose as *de-novo* germline events or represent previously unidentified evolutionary events. In addition, comparison of germline events between the two individuals (D1 and D2), demonstrated that approximately 49% (D1) and 51% (D2) of them were common for both individuals, and among them, 30% (D1) and 31% (D2) were those that were not found in NCBI databases (Supplementary Data S4, S6). Because these two individuals are assumed to be unrelated, it is likely that at least these common novel MMBIR mutations also exist across human populations.

Together, we have identified novel events that were present in human populations, and we propose that both novel and previously identified events were formed through an MMBIR-like mechanism.

### MMBIR mutations arise in a non-small cell lung tumor

To assess whether MMBIR events arise in tumor cells, we first analyzed a lung adenocarcinoma case because it was recently demonstrated that this type of cancer frequently accumulates single-stranded DNA (ssDNA) and also likely uses BIR for DSB repair (Sakofsky et al., 2019; S. Wang, Jia, He, & Liu, 2018), which can enable MMBIR. We extracted genomic DNA and performed WGS on a stage IB lung adenocarcinoma tumor and matched non-tumor lung tissue samples obtained from a 43-year-old patient with a smoking history. We then analyzed the tumor (BO13-UIBB-1654) and non-tumor (BO14-UIBB-1654) derived reads with MMBSearch using identical parameters for both samples and searched for insertions that were a minimum of 10 bp (See Methods for details). To accommodate tumor heterogeneity and enable detection of sub-clonal MMBIR mutations that may exist at very low levels in the sample, we set clustering parameters to a minimum of 3 reads per cluster. With these parameters, and excluding germline calls, we observed that tumor-specific MMBIR calls were 2.3-fold more abundant compared to non-tumor when normalized to the number of sequencing reads for each sample (Figure 3A, B; Supplementary Data S8, S9). We next manually inspected all (414 total) tumor-specific MMBIR calls on chromosome 1 (Figure 3C; Supplementary Data S10). Of the 414 mutations called, 341 (82%) clearly matched the classic MMBIR (CLSC) pattern (Sakofsky et al., 2015). This high level of MMBIR events is in stark contrast to the low number of events found on chromosome 1 of one of the fibroblast clones identified in individual D2 when analyzed with the same MMBSearch parameters (Supplementary text; Supplementary Figure 7A; Supplementary Data S10). The remaining 73 (18%) calls represented noise, which included alterations of microsatellites, undeterminable calls due to poor identity between the insertion and template, and erroneous calls. As an additional control, we isolated genomic DNA from three expanded clonal cultures of lung fibroblast cells and performed WGS using the same library preparation and sequencing method used for the lung tumor samples (see Methods and Supplemental Text). When analyzed with the MMBSearch tool using identical parameters, we observed that only 3% to 5% of MMBIR calls per clone on chromosome 1 matched the MMBIR signature (CLSC) (Supplementary Figure 7B), which was much lower than the level of observed MMBIR events found in the lung tumor. We thus conclude that MMBIR events are highly prevalent in this tumor sample.

**Figure 3.**
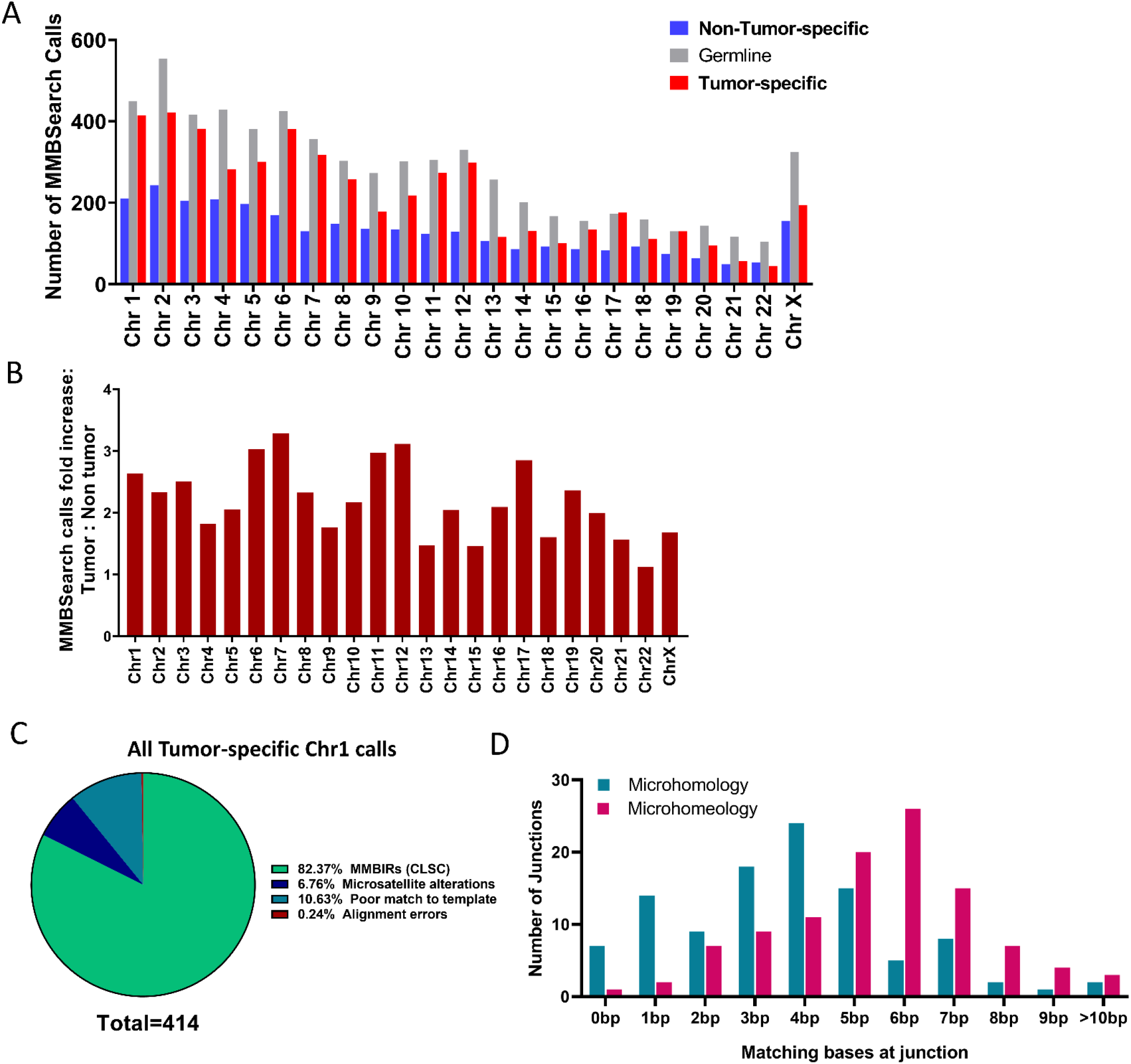
MMBIR events accumulate *de novo* in Lung adenocarcinoma. **(A)** The number of MMBSearch calls per chromosome for lung tumor (BO13-UIBB-1654) and matched non-tumor (BO14-UIBB-1654) tissue samples. **(B)** Fold-increase of MMBSearch calls in tumor-specific calls compared to non-tumor-specific calls (for the counts shown in **A**) normalized to the total number of reads for each individual sample. **(C)** Analysis of all MMBSearch calls on Chromosome 1 in lung tumor from **A**(n=414). Parameters for both samples were ≥3 reads per cluster, insertions that are ≥10 bp in length, and ≥80% identity between template and insertion. **(D)** Microhomology and microhomeology distribution for lung tumor-specific MMBIR event junctions (n=105) from Chromosome1 (dataset shown in **A**(Chromosome 1) and **C**).

Next, to determine the amount of microhomology that mediated the MMBIR events we identified, we analyzed junctions of the first 105 MMBIR events on chromosome 1 by position to determine the length of microhomology used and tolerance for mismatches. We found that 80% of MMBIR event junctions included microhomology sequences of between 2 and 13 bp (average 3.7 bp), and 97% of junctions contained between 2 and 17 bp of microhomeology, with the average length increasing to 5.5 bp (Figure 3D, Supplementary Data S10). Importantly, junctions with 0, 1, and 2 bp of microhomology “acquired” the most additional bases of microhomeology, averaging 3, 2.9, and 2.6 bp gained respectively. Therefore, we propose that practically all events were mediated by microhomology or microhomeology. Moreover, our data suggest that more than 1 bp of total homology was used to initiate these events. Thus, all these events represent authentic MMBIR and were frequent in this cancer sample.

### MMBIR is an initiator of genetic instability cascades

Though the majority of MMBIR events detected were sub-clonal, we hypothesized that MMBIR events with higher clonality might represent initial genomic instability in the tumor, or instability that conferred a selective advantage later in tumor development. By focusing on MMBIR events called by MMBSearch with more than 3 non-duplicate reads, we identified two events, J2 and J6, that were detected by 14 and 11 reads respectively (Figure 4; Supplementary Data S11). Importantly, in addition to an initial MMBIR insertion copied from a nearby template, both of these junctions also included a second region copied from a distant template, resulting in chromosomal rearrangements (Figure 4Ai, iii, Bi, iii). Using BLAST, we confirmed the locations of the secondary templates and designed primers to span the rearrangements (Supplementary Data S12). We confirmed the rearrangements first by PCR and second by Sanger sequencing of the amplified fragments to verify their exact structure (Figure 4Aii, Bii and Figure 5C).

**Figure 4.**
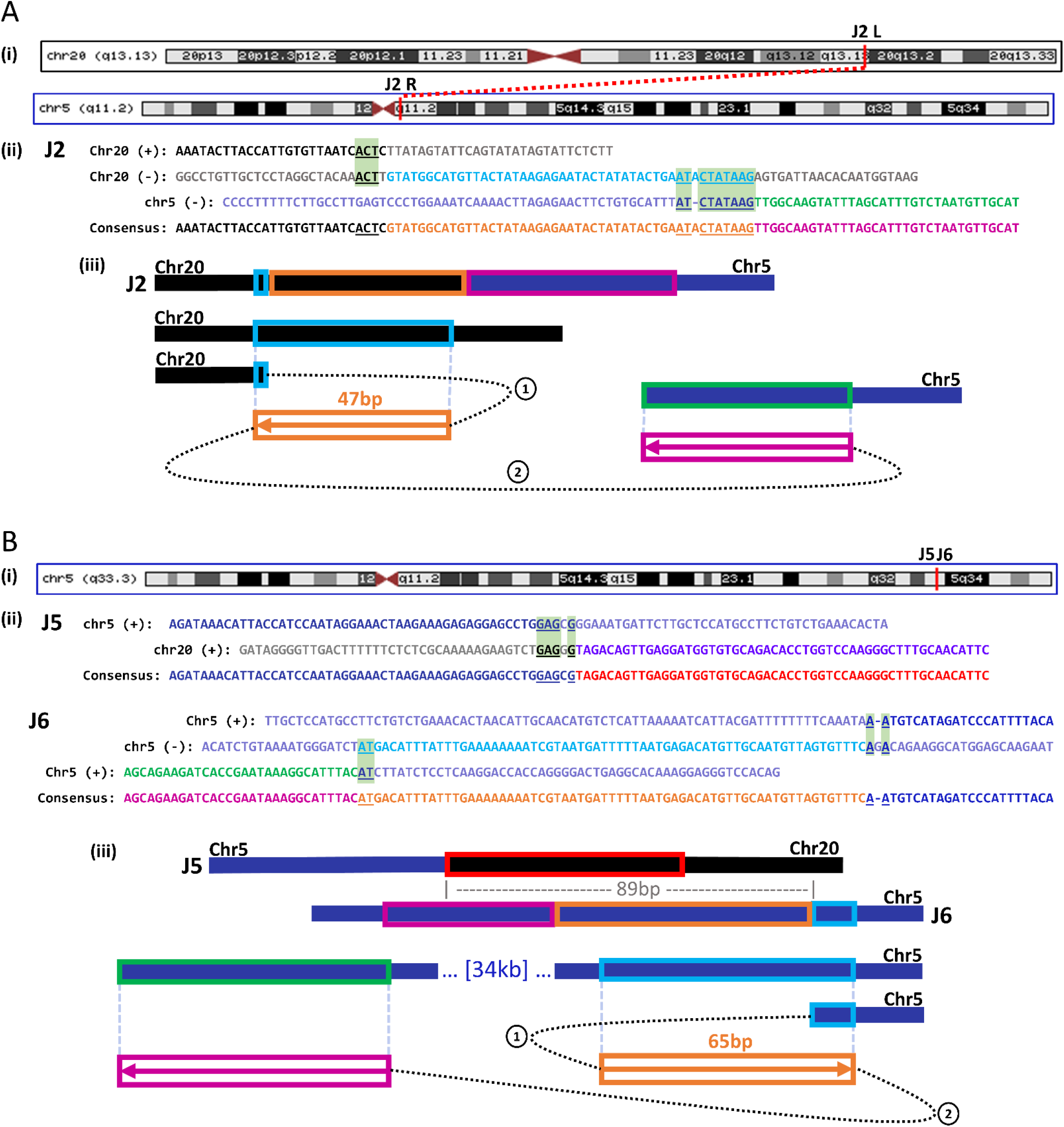
MMBIR initiates genomic rearrangements. **(A) (i)** MMBIR event **J2** found on q13.13 band of Chromosome 20 (Chr20) that is fused to the q11.2 band of Chromosome 5 (Chr5) (red vertical lines indicate positions). **(ii)** Alignment of J2 consensus to all reference templates. Insertion consists of 2 parts (orange and pink text) copied from 2 corresponding templates (light blue and green respectively). Chromosome 20 sequence is shown in black text, and Chromosome 5 sequence is shown in navy. Microhomology used in each template switching event is underlined and highlighted in light green boxes. **(iii)** Schematic illustrating template switching events that lead to the formation of J2. Circled numbers indicate order of template switches. Boxed arrows indicate direction of synthesis, numbers above boxes indicate length of synthesis, and colors correspond to those in **i. (B) (i)** MMBIR events **J5** and **J6** found on q33.3 band of Chromosome 5. **(ii)** Alignment of the J5 rearrangement junction consensus to its reference components on Chromosome 5 and Chromosome 20. Microhomology at J5 is underlined and highlighted in light green boxes. Alignment of event J6 consensus to all reference templates. All labeling similar to **Ai**. **(iii)** Schematic illustrating the structure of J5, its proximity to J6 (89bp), and the two template switching events that lead to the formation of J6. Circled numbers indicate order of template switches. Boxed arrows indicate direction of synthesis, numbers above boxes indicate length of synthesis, and colors correspond to those in **i**.

**Figure 5.**
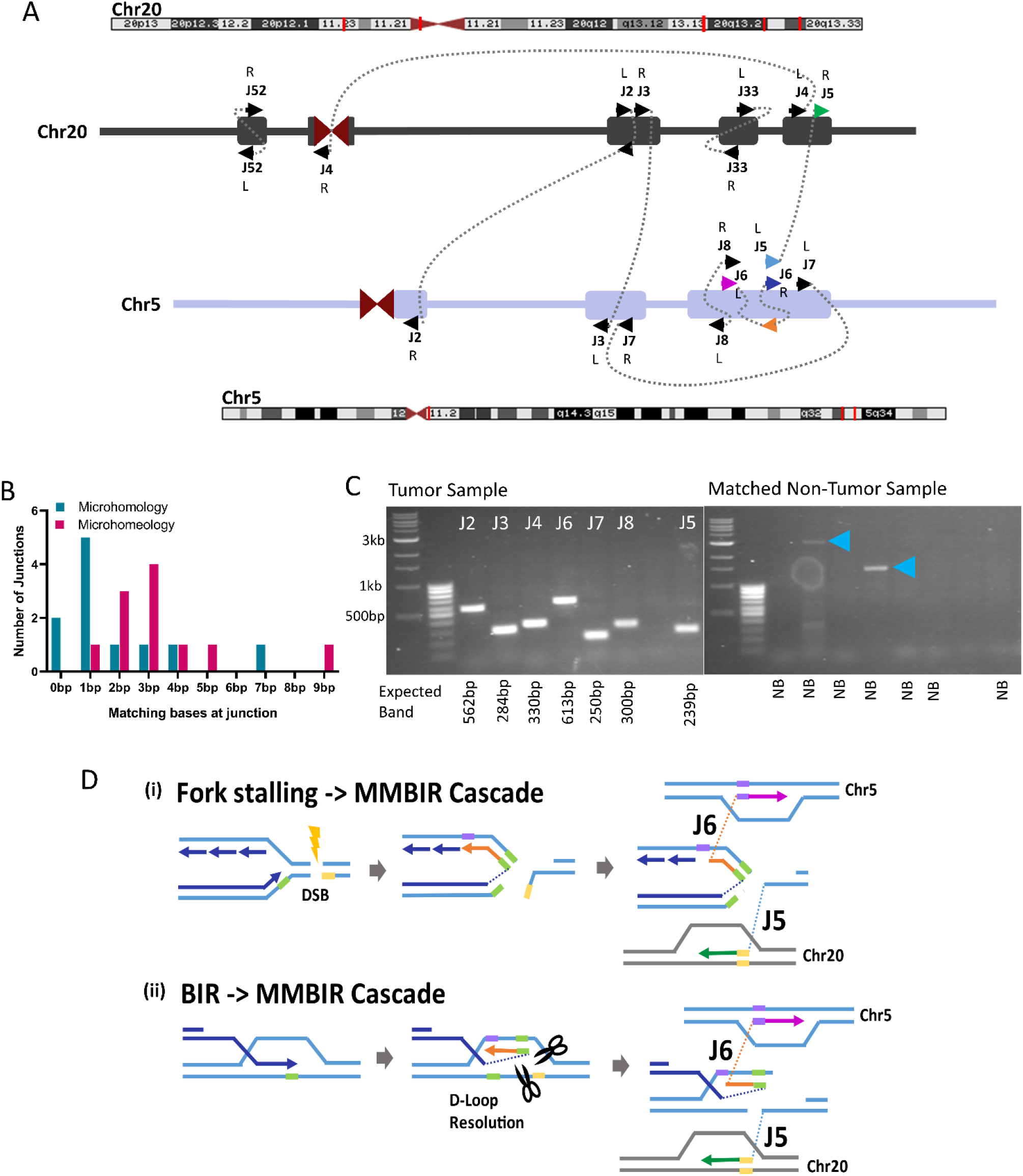
Sub-clonal MMBIR events associated with multiple additional breakpoints on Chromosomes 5 and 20. **(A)** Schematic of all junctions found on Chromosomes 5 and 20. Chromosome positions (red lines on banding diagram) correspond to each cluster of junctions (Rectangles on schematic). L or R indicates Left or Right side of junction with respect to the 5’ to 3’ reference genome. Arrows points indicate forward (to right) or reverse (to left) strands. Arrow colors correspond to the model shown in **D**. **(B)** Distribution of microhomology and microhomeology at all junctions shown in **A**(See Supplementary table S11 for individual junction counts). **(C)** PCR confirmation of MMBIR-related rearrangement junctions shown in **A**. Primers were designed to anneal at positions 100-300bp from the rearrangement junctions and expected band sizes are listed below each lane. Blue arrowheads indicate non-specific bands in non-tumor control reactions. **(D)** Two Models for the formation of junctions J5 and J6 shown in **A**: **(i)** replication fork stalling at a damage site (DSB) leads to template switching of the leading strand to the lagging strand template, and subsequent template switching to distant site on Chromosome 5, while a second broken end invades Chromosome 20. **(ii)** During BIR, the leading strand template switches to the opposite strand within the D-loop structure. D-loop resolution then yields two broken DNA ends that invade different regions on Chromosome 5 and Chromosome 20. Both scenarios lead to genomic rearrangements initiated at an MMBIR event.

The J2 MMBIR event created a fusion between chromosomes 5 and 20 (Figure 4Ai; Supplementary Data S11). Analysis of the junctions suggested that this event likely initiated on chromosome 20, copied 47 bp from a nearby template, and subsequently switched to chromosome 5, leading to a translocation (Figure 4 Aiii). The first junction of this MMBIR mutation did not contain a matching base at the 3’ end itself, but 3 bp directly adjacent to it could act as microhomeology to mediate the template switch. The right junction that mediated the translocation contained 7 matching bases beginning from the 3’ end and gained 2 additional bases when allowing for microhomeology (Figure 4Aii). Importantly, the junction located on chromosome 20 intersects an intron separating the functional exons of the ADNP gene, a transcription factor that is part of the SWI/SNF chromatin remodeling complex (Supplementary Data S11). Mutations in this gene have been identified as a cause of neurodevelopmental disorders such as autism (Helsmoortel et al., 2014)and ADNP has been identified as a potential tumor suppressor in breast and colorectal cancers (Blaj et al., 2017; Rangel et al., 2017). Since this MMBIR event led to chromosomal fusion event between ADNP intron and chromosome 5, it is likely that the ADNP protein will be affected.

The J6 MMBIR event created a chromosomal rearrangement that fused two regions of chromosome 5 located 34 kb away from one another (Figure 4Bi; Supplementary Data S11)). Analysis of the junctions suggested that this event likely initiated from the region telomere-proximal to the fusion, copied 65 bp from a nearby template, and then switched to the centromere-proximal junction (Figure 4Biii). The centromere-proximal junction contained 2 bp of microhomology that could be extended to 3 bp of microhomeology if 1 gap is allowed. The telomere-proximal junction only contained 1 bp of microhomology, and likewise 2 bp of microhomeology (Figure 4Bii J6). Similar to the first MMBIR mutation event, this event results in a fusion between two distinct introns of the EBF1 gene, which encodes a transcription factor necessary for B-cell progenitor differentiation. Mutations in EBF1 are common in acute lymphoblastic leukemia (Heinrichs & Look, 2007) (Supplementary Data S11). Furthermore, heterozygosity of this gene is associated with increased DNA damage and decreased homologous recombination through decreased Rad51 expression and decreased apoptosis in mice (Heinrichs & Look, 2007).

The higher read evidence for these two particular MMBIR mutation events, as compared to other MMBIR mutation calls (Supplementary Data S10), suggests a higher level of clonality of these two mutations in the tumor. Because these events involved chromosomes 5 and 20, we asked whether there were additional junctions nearby on these two chromosomes with similar levels of read evidence that might be related to these MMBIR events. By searching the clusters of reads generated by MMBSearch, we found 7 additional breakpoint junctions involving chromosomes 5 and 20 that were detected between 9 and 15 reads each and that were confirmed by PCR (Figure 5A, C; Supplementary Data S11, S13). All of these similarly evidenced junctions also contained varying lengths of microhomology (Figure 5B; Supplementary Data S11, S13). While these breakpoints are not sufficiently clonal to produce a change in ploidy, the increased levels of reads associated with them implies a sub-clonal increase in copy number (Supplementary Figure 8A, B). Analysis of clusters also revealed that the cluster containing MMBIR event J6 contained a second junction, J5. The first breakpoint for J6 and the J5 breakpoint were less than 100 bp from each other (Figure 4Bii, iii), and this proximity strongly suggests a single breakage event. Importantly, this implies that the MMBIR event found at J6 was the initiating template switching event that led to discoordination of two ends from a single break, resulting in two distinct rearrangements at a single locus. Because the J6 MMBIR event replaces the majority of its template, we can speculate that it was copied from either the lagging strand in the context of a stalled replication fork (Figure 5Di), or from the top strand within the BIR D-loop (Figure 5Dii), before template switching to a more distal template, while the J5 junction forms as the result of template switching initiated by a second broken end (Figure 5Di, ii).

In addition to events that did not produce a change of ploidy, Chromosomes 5 and 20 in the tumor also displayed 3n regions (Supplementary Figure 8A) implying that the breakpoints of those 3n regions were fully clonal, and likely existed starting from the point the tumor was established. We investigated these fully clonal rearrangement events and found a similar pattern of rearrangements and coincident breakpoints, one of which again contained an MMBIR event on one of two coincident junctions just 63 bp away from each other (Supplementary Data S11, Junctions J14 and J15).

We propose that the coincident breakpoints observed at different levels of clonality represent the initial breaks from which all other observed rearrangements with similar levels of clonality proceeded. Further, the presence of MMBIR events at these coincident junctions implicate MMBIR as an initiator of a cascade of genomic rearrangements, which culminate to chromothripsis. Thus, by searching for the initiators of MMBIR with MMBSearch we were able to unravel entire complex rearrangements that are similar to the MMBIR events described in (Carvalho et al., 2013; Carvalho et al., 2011; Cortes-Ciriano et al., 2020; Liu et al., 2011) in connection to neurological diseases and chromothripsis.

### Analysis of TCGA cancers reveals increased levels of MMBIR mutations

To determine whether MMBIR events arise at a high frequency across other types of cancer, we analyzed tumor samples and their matched non-tumor controls obtained from The Cancer Genome Atlas (TCGA) and dbGaP. For this analysis, we extracted all unmapped reads for each sample and analyzed them with MMBSearch using parameters identical to those used for the stage IB lung adenocarcinoma sample (see Methods for details). The detailed analysis of a lung tumor sample obtained from dbGaP (see Methods) produced results similar to those obtained from the stage IB lung adenocarcinoma sample. In particular, after excluding germline MMBIR events, we observed 2.6-fold more tumor-specific MMBIR calls compared to matching non-tumor samples (Figure 6A, B; Supplementary Data S14). Manual inspection of all chromosome 1 tumor-specific calls revealed that 85% matched the classic MMBIR (CLSC) pattern (Figure 6C). In addition, 95% of analyzed MMBIR junctions contained 2 or more bases of microhomology or microhomeology (Figure 6D; Supplementary Data S15, S16). We further confirmed that some of the MMBIR events were complex, as they resulted from several consecutive template switching events mediated by microhomologies, and some of these also resulted in chromosome fusions (Figure 6E, F; Supplementary Data S17) (See Supplementary text for full description of these events). Differently from the stage IB Lung cancer (Figures 3–5), in this cancer sample all identified CGR events initiated by MMBIR had low clonality in the sample (Supplementary Data S17).

**Figure 6.**
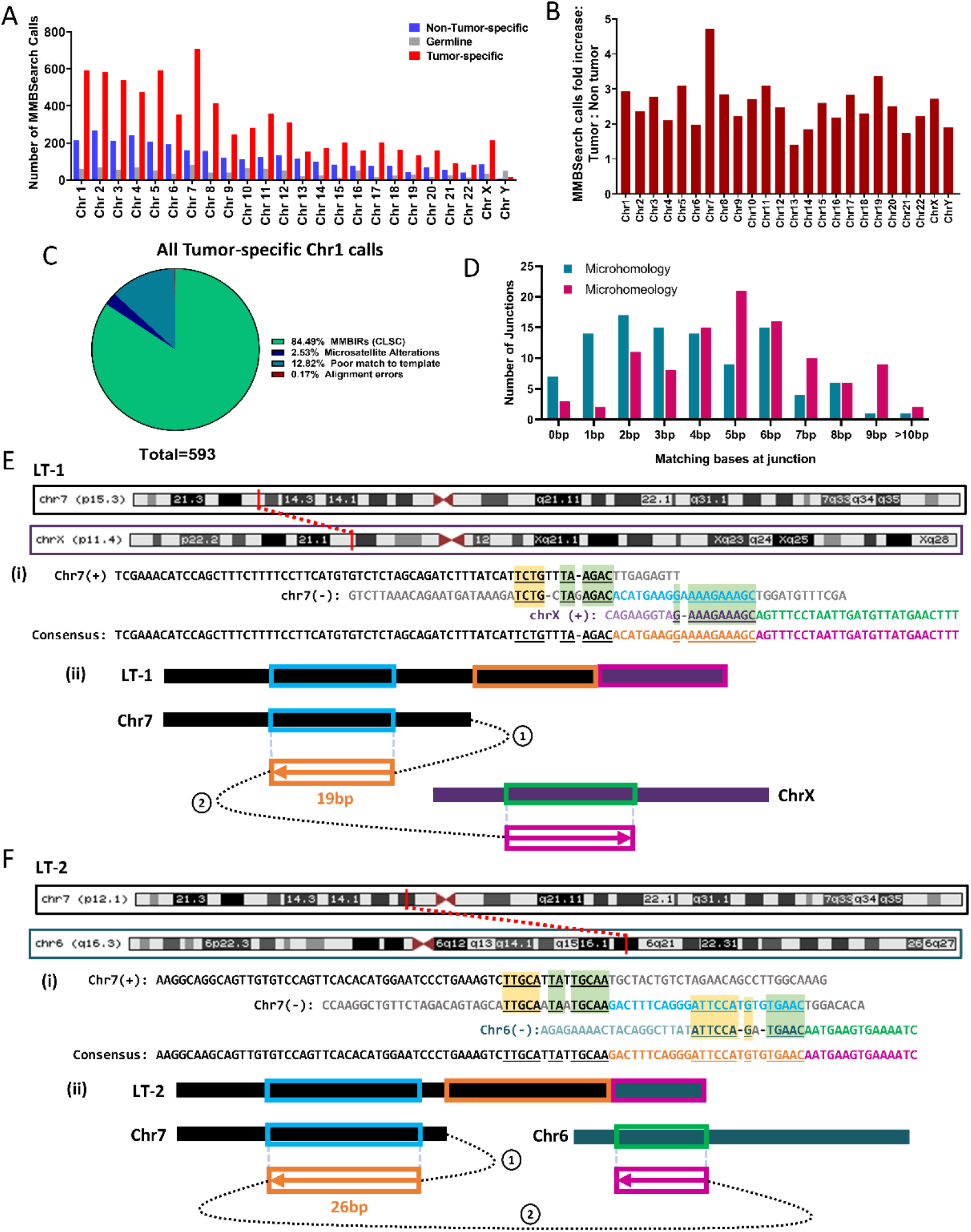
MMBIR events accumulate in a Lung adenocarcinoma case from dbGaP (accession ID: phs000488.v2.p1). **(A)** MMBSearch calls per chromosome for lung tumor (SRA run ID: SRR556475) and matched non-tumor (SRA run ID: SRR551334) tissue samples. **(B)** Fold-increase of MMBSearch calls in tumor specific-calls compared to non-tumor-specific calls (for the samples shown in **A**) normalized to the total number of reads for each individual sample. **(C)** Analysis of all MMBSearch calls on Chromosome 1 for lung tumor (n=593). Events were detected using identical parameters to those used in analysis for Figure 3C. **(D)** Microhomology and microhomeology distribution for MMBIR event junctions from Chromosome 1 (n=103) of the same lung tumor sample. (**E)** MMBIR event LT-1 found on p15.3 band of Chromosome 7 that is fused with the p11.4 band of Chromosome X. **(i)** Alignment of event LT-1 consensus to all refence templates. Insertion consists of 2 parts (orange and pink text) copied from 2 corresponding templates (light blue and green respectively). Chromosome 7 sequence is shown in black text, and Chromosome X sequence is shown in purple. Microhomology used in each template switching event is underlined and highlighted in light green boxes, and additional microhomology is highlighted in yellow boxes. **(ii)** Schematic illustrating template switching events that lead to the formation of LT-1. Circled numbers indicate order of template switches, Boxed arrows indicate direction of synthesis, numbers above boxes indicate length of synthesis, and colors correspond to those in **i**. **(F)** MMBIR event LT-2 found on p12.1 band of Chromosome 7 that is fused with the q16.3 band of Chromosome 6. **(i)** Alignment of event LT-2 consensus to all refence templates. Chromosome 7 sequence is shown in black text, and Chromosome 6 sequence is shown in teal. Other text colors and highlights similar to those in **E i**. **(ii)** Schematic illustrating template switching events that lead to the formation of LT-2. Circled numbers indicate order of template switches, Boxed arrows indicate direction of synthesis, numbers below boxes indicate length of synthesis, and colors correspond to those in **i**.

We next determined the number of MMBIR events in a total of 71 tumor and matching non-tumor samples obtained from TCGA (Supplementary Data S18). To accommodate varying depths of sequencing coverage across many samples, we normalized the number of MMBIR events called to the total number of reads for each sample (Supplementary Data S19). Across the 7 cancer types analyzed, tumors from ovarian cancer, lung adenocarcinoma, colon adenocarcinoma, and endocervical adenocarcinoma sites had significant increases in the number of calls made by MMBSearch in the tumor samples compared to their matched non-tumor counterparts (Figure 7A; Supplementary Data S19). Breast invasive carcinoma tumors did not show a significant increase overall, but some cases displayed exceptionally high levels of MMBIR events in the tumor samples compared to non-tumor (4.7-to 23-fold increase). This was also true of some lung adenocarcinoma tumors, which had fold-increases of 3.2-to 8.6-fold (Figure 7B). These data suggest that MMBIR events arise in tumors from multiple primary sites and accumulate in tumor cells, resulting in elevated levels of MMBIR events in tumors.

**Figure 7.**
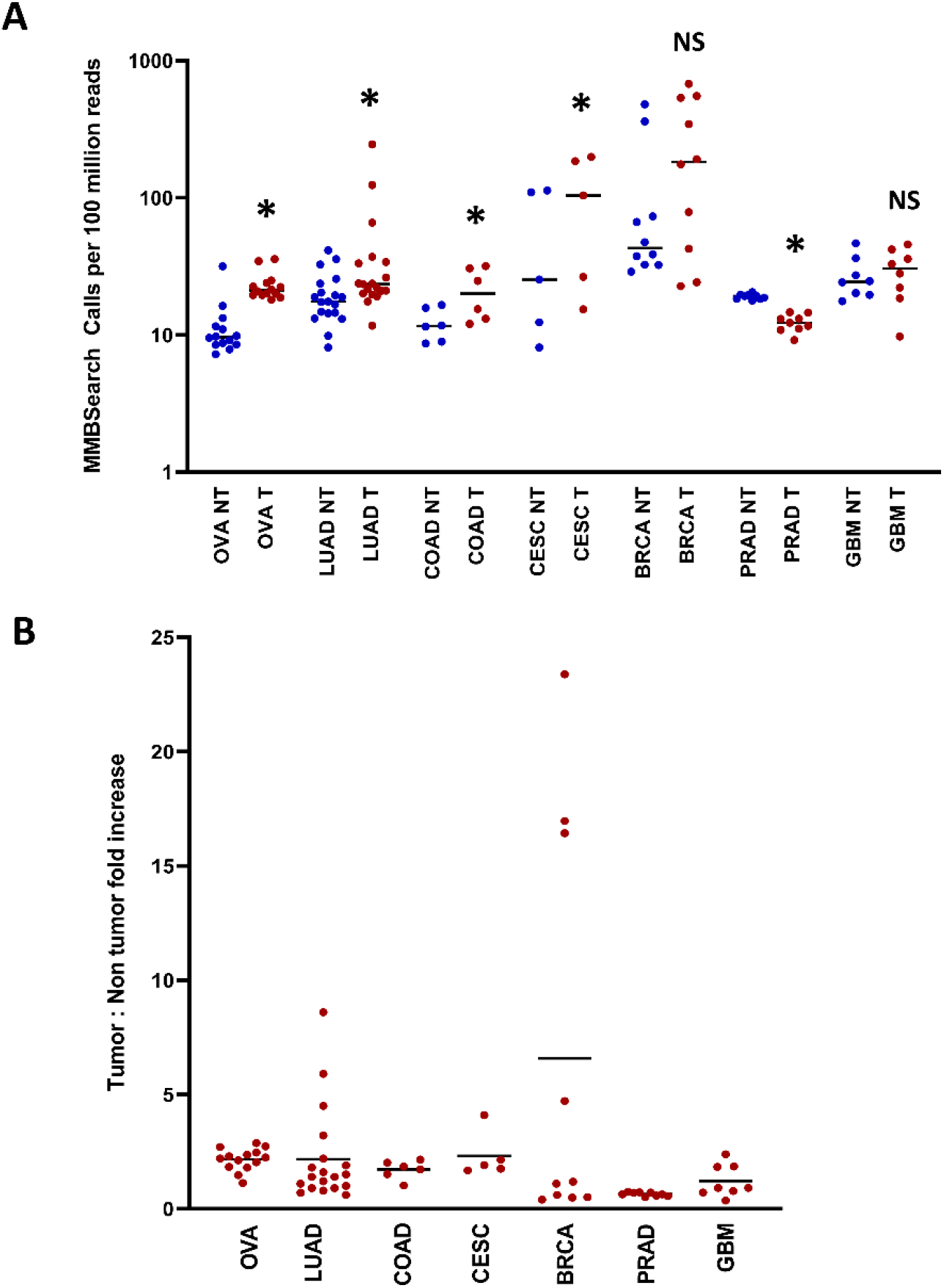
The level of MMBIR events is increased in tumors from multiple primary tumor sites. **(A)** Number of MMBSearch calls per 100 million reads quantified for paired tumor (red) and non-tumor (blue) samples. WGS data was obtained from 71 total TCGA cancer cases. Asterisks indicate statistically significant difference in the level of MMBIR events between tumor and non-tumor for each cancer site (p-value<0.05) calculated by ratio paired t-test. NS = no significant difference. Cancer primary sites are as follows: Ovarian cancer (OVA) (n=14), lung adenocarcinoma (LUAD) (n=19), colon adenocarcinoma (COAD) (n=6), cervical squamous cell carcinoma and endocervical adenocarcinoma (CESC) (n=5), breast invasive carcinoma (BRCA) (n=10), prostate adenocarcinoma (PRAD) (n=9), and glioblastoma multiforme (GBM) (n=8). Black lines indicate median values for each set. **(B)** Fold-increase in the amount of tumor vs. non-tumor MMBSearch calls shown in **A**. Each dot represents one cancer case (tumor compared to matched non-tumor). Black lines indicate mean fold-increase.

## Discussion

In this study, we employed our novel MMBSearch software tool to identify MMBIR events based on the signature of MMBIR with unprecedented resolution and irrespective of whether they produced a CGR or CNV. This approach allowed us to determine the frequency of MMBIR events in different cell contexts, which uncovered highly disparate frequencies in normal human cells compared to cancer cells. While MMBIR events were rare in somatic fibroblasts, they were frequent in cancers. Moreover, MMBIR was detected only as germline events in fibroblasts, whereas they accumulated over time in cancer cells and contributed to genetic instability.

Based on our data, we propose a model of MMBIR and its various genomic outcomes (Figure 8). In this model, as a result of fork stalling or induction of a double strand break (Figure 8i), a 3’ single-strand DNA (ssDNA) end formed by dissociation of a nascent strand from its template during S-phase replication, or by 5’-to-3’ resection of DSB ends, can anneal to exposed ssDNA at a site of microhomology, or invade double-strand DNA (dsDNA) using microhomology to prime synthesis (Figure 8ii). Subsequent template switching events can either return to the original template or anneal elsewhere, resulting in a more complex rearrangement (Figure 8iii). More localized MMBIR events that do not disrupt genes may accumulate as germline variants over the course of evolution, while more complex MMBIR events, especially those that disrupt genes or lead to CGRs, are more likely to appear only in pathogenic contexts (Figure 8iv).

**Figure 8.**
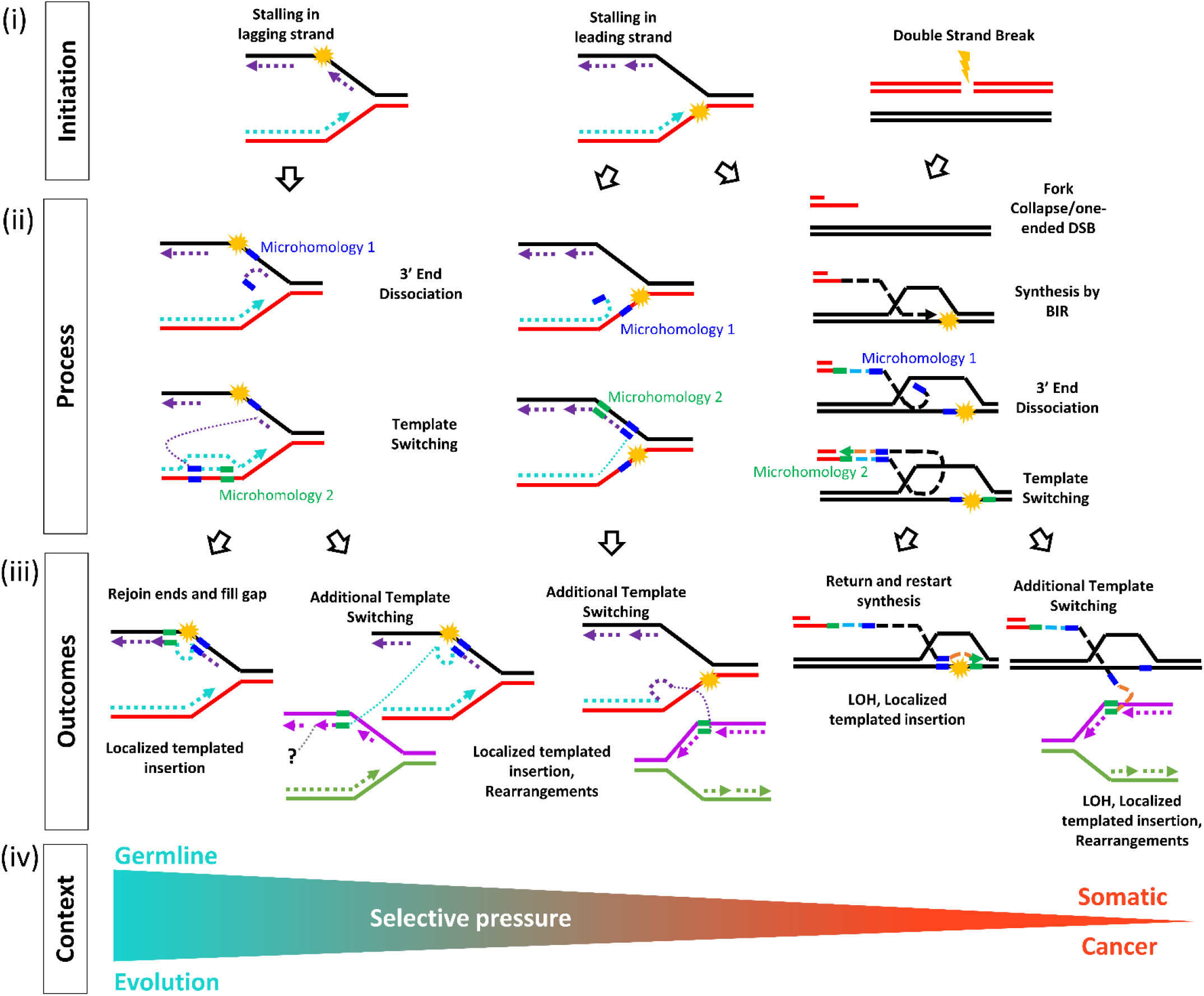
Model explaining formation of MMBIR and its various genomic outcomes. **(i) Initation:** Stalling of leading, lagging strand, or induction of a double strand break lead to replication problems or induction of BIR. **(ii) Process:**3’ single-strand (ss) DNA ends are formed by dissociation of a nascent strand from its template during S-phase replication or by 5’ to 3’ resection of DSB ends. Dissociated or resected 3’-ssDNA anneals at microhomology to nearby exposed ssDNA or invades nearby dsDNA at microhomology to prime DNA synthesis. **(iii) Outcomes:** Additional rounds of template switching, or invasion mediated by microhomology can lead to either a return to the original template or more complex rearrangements. **(iv) Context:** MMBIR events that do not disrupt genes are more likely to accumulate as germline variants over the course of evolution. More complex MMBIR events, especially those that disrupt genes or lead to gross chromosomal rearrangements, may only appear in pathogenic contexts.

### Initiation of MMBIR involves more microhomology than previously demonstrated

Our analysis determined that template switching associated with MMBIR events involved microhomologies, which is consistent with our proposed molecular mechanism (Figure 8). Previous attempts to determine the microhomology requirements for chromothripsis (Cortes-Ciriano et al., 2020) suffered from collectively analyzed events originating from different repair mechanisms, making it impossible to determine the microhomology requirements for MMBIR specifically. Our analysis of MMBIR events in lung adenocarcinomas demonstrated that nearly all MMBIR events involved a significant amount of microhomology. We further report that these microhomologies can often be extended by allowing gaps or mismatches. In particular, we observed that addition of tolerance for a single unmatched base (gap or mismatch) universally extended microhomology at all junctions, but especially at those that were previously recorded as having 0 to 1 bp of microhomology. The implications of this are two-fold. First, it suggests that the actual amount of homology typically used in MMBIR is higher than previously appreciated. Second, it indicates that the annealing (pairing) between strands initiating MMBIR template switching does not have to be perfect. In fact, based on our findings, even the 3’-most base does not have to be matched to the template to prime synthesis.

The concept of imperfect microhomology, or microhomeology, that we used in this work is different from what has been described in previous studies that attempted to account for microhomology containing mismatches (Liu et al., 2017). The authors attempted to define microhomeology by gapped alignment of junction-flanking sequences, which demonstrated that microhomology could be imperfect, but did not specify cases with a mismatch at the 3’ end implicated in priming the synthesis. In our analysis, in which we identified unmatched bases beginning from the 3’ end, we were able to demonstrate that MMBIR can be initiated even when the bases closest to the 3’ end are mismatched. The observation that MMBIR can utilize microhomeologous sequences allows us to speculate that MMBIR is unlikely to be carried out by polymerase(s) that conduct S-phase DNA synthesis, which possess efficient exonuclease function that would be expected to remove mismatched bases before extending. Translesion polymerases are likely candidates for this function. Indeed, in yeast, we previously demonstrated that polymerase ζ is responsible for MMBIR synthesis (Sakofsky et al., 2015).

Another candidate polymerase to initiate synthesis from imperfect primers in mammalian cells is Polθ, which is known to form insertions and deletions by initiating DNA synthesis from primers annealed at microhomologies in theta-mediated end-joining (TMEJ) (Beagan & McVey, 2016; Schimmel, Kool, van Schendel, & Tijsterman, 2017). It has been recently shown that, during DSB repair, TMEJ favors annealing of a broken 3’ end at microhomologies most proximal to DSBs (Carvajal-Garcia et al., 2020). Furthermore, Polθ has been shown to aid in repair of collapsed replication forks (Z. Wang et al., 2019)which can serve as substrates for the initiation of MMBIR. It is possible that Polθ-mediated TMEJ may represent at least a subset of the MMBIR events that we uncovered in this study. Because the literature regarding TMEJ has focused on shorter, less complex insertions than we associate here with MMBIR, we are unable to discern whether any of the events we describe as MMBIR should be attributed to TMEJ.

### MMBSearch as a tool to detect MMBIR

In this study, we used our new software tool to detect the initial, and sometimes multiple template switches of MMBIR events. Consistent with our initial hypothesis, many of these events were discarded by conventional BWA mapping methods, where they were either unmapped or among clipped reads, depending on the parameters of mapping. We previously developed a software tool for direct detection of MMBIR events in the yeast genome (Segar, Sakofsky, Malkova, & Liu, 2015), but that approach lacked the efficiency and flexibility required to query the larger datasets of human genomes. MMBSearch overcomes these limitations through a novel parallel computing approach that makes it suitable to identify MMBIR signatures efficiently within human genome datasets. MMBSearch accurately calls MMBIR events. Its sensitivity is especially high for identifying insertions without replacement, and somewhat lower for insertions with replacement, which likely results from the leveraging of reads that differ greatly from the reference.

The results from the present study indicate that MMBSearch should be automated to enable identification of subsequent template switches to distant and nearby templates. This currently requires manual curation of events to identify additional templates for highly complex MMBIR events. Automation of this process would contribute to our understanding of the mechanism that generates complex MMBIR patterns like those discovered in this study. In addition, automation for determining the amount of microhomology and microhomeology associated with MMBIR represents another important goal.

### MMBIR: a driver of cancer evolution?

Our results suggest that normal human cells differ from cancer cells in their propensity for MMBIR. In the genomes of normal skin fibroblasts from two individuals, MMBIR signatures were only present as germline variants, which differentiates MMBIR events from all other mutations, including base substitutions and frameshifts that have been observed to accumulate with age in these cells (Saini et al., 2016). Some of the MMBIR events were also found as a subset of germline events across human populations from other studies, or were shared between the two individuals, implying that these events may have accumulated over the course of human evolution. While these germline MMBIR mutations could be complex, having copied from up to 5 templates to form a single event, they generally did not lead to CGRs. This is expected, as the selective pressure imposed during evolution would likely eliminate detrimental MMBIR events from the genome, except in some rare cases of congenital diseases.

In tumors, across a variety of TCGA cohorts, our results indicate that MMBIR events accumulate *de novo*. Most accumulated MMBIR events were sub-clonal, but in our detailed analysis of lung adenocarcinomas, we specifically found some rare MMBIR events tied to GCR junctions that had expanded in the tumor. Importantly, we observed these expanded MMBIR events disrupting DNA repair (EBF1) and chromatin remodeling (ADNP) genes. Based on this observation, we propose that MMBIR may play a critical role in cancer initiation, progression, and innate drug resistance by mutating tumor suppressor or oncogenes. For tumors treated with chemo- or radiotherapy, in which DSBs are induced *en masse*, MMBIR may play a key role in adaptive response and disease relapse. A more comprehensive understanding of how the mutational landscape of tumor cells influences the propensity for MMBIR is needed to determine why some rearrangements may be preferentially selected in certain genetic contexts.

In matched-normal tissues from donors of tumors where MMBIR events accumulated, we often found high levels of MMBIR. Although there was a general trend of increased MMBIR load in tumor compared to matched normal samples, the relatively high MMBIR load in normal tissue suggests that MMBIR events may still accumulate in some tissues of cancer patients, even if our analysis of clonal fibroblast lineages of healthy individuals did not uncover accumulation of MMBIR events. This may have been influenced by the mutational landscape of the donors of fibroblast versus donors of tumor tissue, or it may indicate that the predisposition of different tissue types to undergo DNA damage and repair can influence the frequency of MMBIR. Extensive analysis of DNA samples from different tissues and across populations of healthy donors would be required to improve our understanding of how and where MMBIR events accumulate. One other possible source that could contribute to the finding of elevated MMBIR events in non-tumor samples is contamination by tumor DNA, particularly if tumor cells are actively being destroyed by treatment or the immune system. Studies have found tumor DNA circulating in the blood as cancer cells are eliminated (Cristiano et al., 2019; Diehl et al., 2008; Sozzi et al., 2003), and identification of tumor-specific mutations from circulating tumor DNA (ctDNA) in the peripheral blood is the basis for cancer diagnostic blood tests (Cohen et al., 2018; Wan et al., 2017). This provides one explanation for why some matched non-tumor samples, particularly whole blood, have a high frequency of MMBIR events, and suggests that future studies to measure MMBIR mutation signature load in blood could be useful as a diagnostic or prognostic biomarker for some types of cancer.

## Methods

### Read preprocessing and the MMBSearch tool

MMBSearch was used with either raw reads files or reads extracted from an alignment (BAM) file. Samtools (version 1.9) was used to extract unmapped reads from BAM files via the alignment flag, while soft-clipped reads and reads with indels were extracted from gapped alignments (those aligned by BWA-MEM (Li, 2013)) by selecting reads containing “I”, “D”, or “S”, in their CIGAR string. PRINSEQ (Schmieder & Edwards, 2011) was used to perform low-complexity filtering via the DUST algorithm (threshold=7). Trimmomatic (version 0.38) (Bolger, Lohse, & Usadel, 2014) was used to perform adapter trimming with the Truseq3 Single-end adapter set and to perform trimming of low quality bases and N bases from read ends using the following parameters: LEADING:3 TRAILING:3 SLIDINGWINDOW:4:15 MINLEN:36.

After preprocessing, MMBSearch aligns reads using BWA-backtrack and an error threshold of 4. MMBSearch then recovers all unmapped reads (Those with more than 4 differences from the reference) and splits them in half. The half-reads are again aligned to the reference by BWA-backtrack with the same error threshold, and the reads with one mapping and one non-mapping half are accepted for clustering. Clusters that meet the user-specified threshold of reads are analyzed by their anchoring side (mapping half-read for each read in the cluster). Clusters that anchor all left, all right, or left and then right are accepted, and consensus sequences are called for the anchored reads. The two consensuses that form from left-then-right clusters are locally aligned to each other to correct any positional errors. The full consensus contigs are then compared back to the reference sequence to determine the sequence of a candidate MMBIR region (insertion or breakpoint). The sequence of the MMBIR region is then used to search for a nearby template (within 100 bp). MMBSearch outputs files containing the consensus contigs, full clusters, and called MMBIR events (a list of insertions, reference locations, and alignments of each MMBIR insertion to its template).

Reads derived from clonal skin fibroblasts (Saini et al., 2016) (NCBI accession: PRJNA336369; dbGaP: phs001182), and reads derived from dbGaP IB lung adenocarcinoma (NCBI accession: PRJNA157939; dbGaP: phs000488) were analyzed with respect to human genome reference version GRCh37/hg19 to maintain positions with previous studies, while all other samples were analyzed with respect to GRCh38/hg38 for posterity.

Fibroblast data sets were analyzed using the following MMBSearch configuration parameters: For the initial analysis of all events, minimum MMBIR event length was set to 10 bp with a minimum 80% identity between the MMBIR event and its reference template. Secondary analysis to find more complex MMBIR events and to eliminate interference from shorter microsatellites was achieved by setting the minimum MMBIR length to 25 bp, with a minimum of 40% identity between the MMBIR event and the template. For both analyses, minimum cluster size was set to 10 reads. Cluster size was reduced to 3 reads for the sample used for comparison to cancer genomes (Supplementary Figure 7). Similarly, for both lung adenocarcinoma datasets and all TCGA datasets, minimum MMBIR event length was set to 10 bp with a minimum identity of 80% between MMBIR event and template. The two lung adenocarcinoma samples analyzed in depth underwent secondary analysis allowing a minimum of 20 bp or 25 bp depending on read length (100 bp and 150 bp respectively) with 40% identity between MMBIR event and template. For all cancer datasets, minimum cluster size was set to 3 reads (All other MMBSearch configuration parameters were left as default unless otherwise specified). The number of reads of evidence for MMBIR events and structural variant (SV) junctions were used to infer clonality by dividing the number of MMBIR or SV event reads by the reported average mappable coverage of the genome. Events with read counts less than 30% of the average mappable coverage were considered to be sub-clonal. Fibroblast and lung adenocarcinoma datasets that were analyzed in-depth were manually inspected to exclude false positive calls and microsatellite calls, and had positions of germline (Fibroblasts) and sub-clonal to clonal (Lung adenocarcinoma) MMBIR events confirmed using BLAT (Kent, 2002) to exclude ambiguous calls and to help resolve more complex multi-templated MMBIR events. BLAST (Altschul, Gish, Miller, Myers, & Lipman, 1990) was used to query NCBI’s Nucleotide collection to search for previously published germline MMBIR insertions.

For manually inspected junctions with resolvable templates, microhomology was counted by comparing the insertion junctions identified by MMBSearch in sequencing reads to the reference-derived complement to the insertion (template) and its flanking sequence. Microhomology was defined as uninterrupted matching bases at the junction. Likewise, the allowance of a single mismatch or gap that interrupts the matching bases at a junction and results in at least 1 additional complementary base is referred to as “microhomeology.” Read aggregate analysis of lung adenocarcinoma samples BO13 and BO14 to determine changes in copy number was performed using CLC genomics workbench 8. Ploidy was inferred by comparing read coverage in regions with apparent copy number changes to estimated average coverage for the genome.

### MMBSearch Sensitivity testing

We generated 993 artificial MMBIR insertions of various sizes (20-50 bp) in GRCh37/hg19 Chromosome 17 at regular intervals, excluding regions of Chromosome 17 that contain N bases. Using ART (version: MountRainier-2016-06-05) (Huang, Li, Myers, & Marth, 2012) we then simulated paired-end 125 bp Illumina reads with an average insert size of 500 bp with a standard deviation of 20, to an average depth of 30x. These reads were analyzed with MMBSearch with search parameters set to a minimum MMBIR length of 6 bp with a minimum of 3 reads to form a cluster. The results output was then analyzed for true positives (TP) by comparing MMBSearch MMBIR calls to the list of artificial MMBIR insertions. Remaining calls or duplicate calls were considered to be false positives (FP). The number of false negatives (FN) was determined by subtracting TP from the total events inserted. Recall was calculated with the following formula: TP/(TP+FN).

### Molecular Biology Methods

Genomic DNA was prepared from tumor and non-tumor tissues frozen in OCT medium following tumor resection using the Qiagen blood and Cell culture DNA Mini Kit (Cat no. 13323). Library preparation was performed for whole genome sequencing using the TruSeq DNA PCR-Free (350) kit (protocol: TruSeq DNA PCR-Free Sample Preparation Guide, Part # 15036187 Rev. A). Sequencing was performed on the Illumina NovaSeq6000 S4 (2×150 bp) sequencing platform. Tumor (BO13-UIBB-1654) tissue was sequenced to an estimated average coverage of 51x, and non-tumor (BO14-UIBB-1654) tissue was sequenced to an estimated average coverage of 65.5x.

GM1604 human fetal lung fibroblast cells were cultured in Dulbecco’s Modified Eagle Medium (DMEM, Life Technologies) supplemented with 10% Fetal Bovine Serum (FBS, Gibco) and 100 Units/mL of Penicillin/Streptomycin solution (Life Technologies). Cells were incubated at 37°C and 5% CO2 in a humidified atmosphere. GM1604 clones were obtained by expanding cell colonies originating from a single cell. Genomic DNA from the expanded human fetal lung fibroblast clones was purified using the Qiagen blood and Cell culture DNA Mini Kit. Library preparation was performed for whole genome sequencing using the TruSeq DNA PCR-Free (350) kit (protocol: TruSeq DNA PCR-Free Sample Preparation Guide, Part # 15036187 Rev. A). Sequencing was performed on the Illumina NovaSeq6000 S4 (2×150 bp) sequencing platform to achieve an estimated average coverage of 26-32x.

MMBIR and associated rearrangement junctions found in Lung tumor (BO13-UIBB-1654) were confirmed by PCR. Forward and reverse primers were designed to the boundaries of each junction (See Supplementary Data S12 for all primers used), and PCR was performed using Phusion High-fidelity DNA Polymerase (NEB Cat no. M0530S). PCR products of junctions containing MMBIR events were sequenced by Sanger sequencing to confirm the presence of the MMBIR insertion within the rearranged allele. All PCRs were also performed with non-tumor DNA from the same patient (BO14-UIBB-1654) to confirm tumor specificity of the rearrangement junctions.

## Data Availability

The MMBSearch tool package and all additional code associated with this study, including the code used to generate artificial datasets, can be downloaded from https://github.com/malkovalab/MMBSearch. The majority of the datasets analyzed in this study are available from the The Cancer Genome Atlas (TCGA) project (https://portal.gdc.cancer.gov/) or the database of genotypes and phenotypes (dbGaP) authorized access portal. Accession IDs for all samples from databases can be found in Supplementary data S1, S2, S14, and S18 as either SRA run IDs or GDC case IDs. Raw sequencing reads from all prepared genomic DNA are available through dbGaP (Study accession ID pending). All data supporting the findings of this study are represented in this article or supplementary materials. Raw output files are available upon reasonable request.

## Acknowledgments

We thank Drs. Dmitry Gordenin, Natalie Saini, and members of the Malkova Lab for helpful comments on this manuscript. We also thank Tara Hicks and Katelyn Morrison for their help with fibroblast and cancer MMBSearch data analysis. This work was supported by NIH grants R35GM127006 and R01CA232425 to AM.

## Supplementary Information

### Supplementary Figures

**Supplementary Figure 1:**
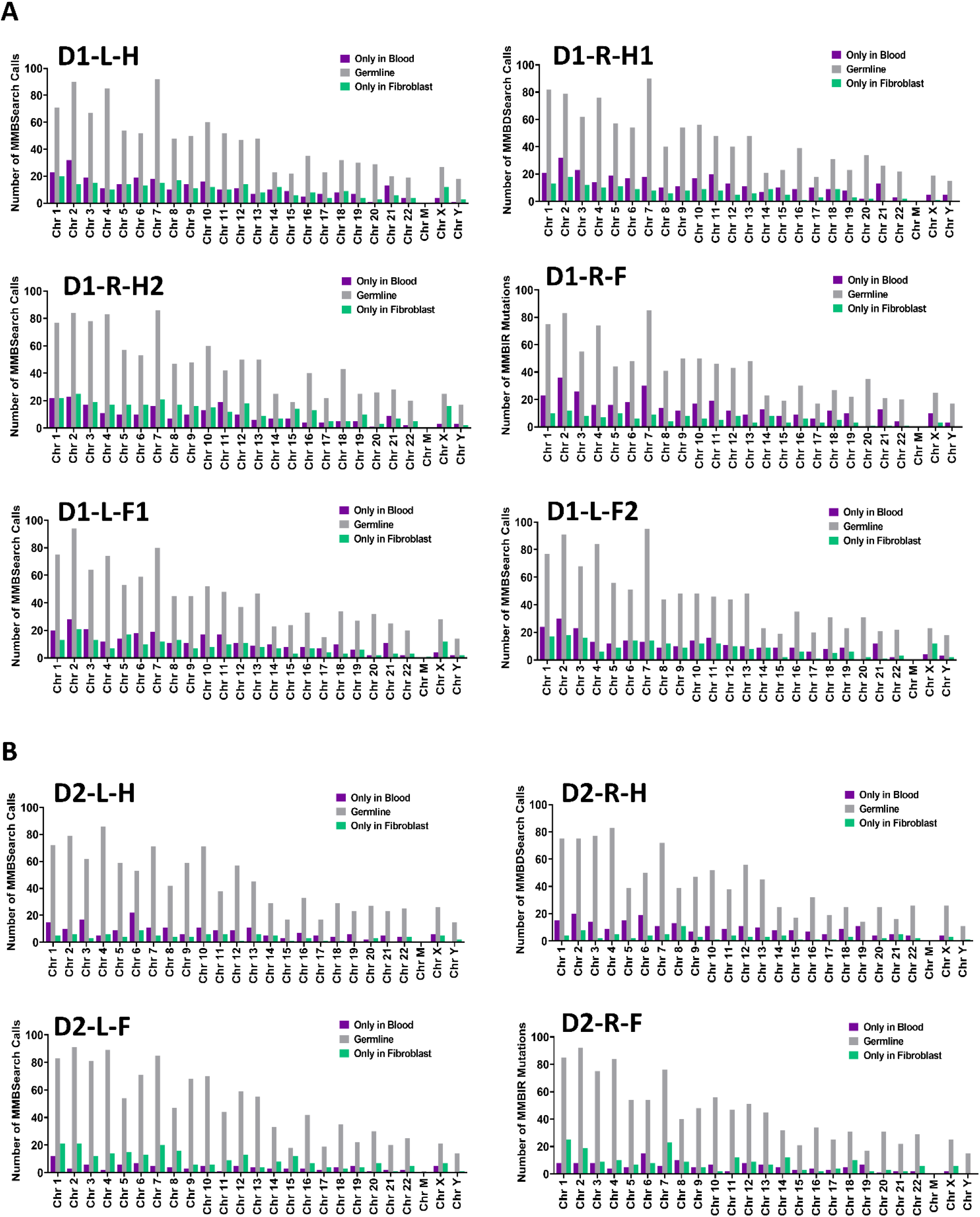
MMBsearch calls for all fibroblast clones. MMBSearch calls (from clusters with ≥10 reads, insertions that are ≥10bp in length, and ≥80% identity between template and insertion) per chromosome for 10 fibroblast cell lineages from the study of ^2^. Germline calls are those found to overlap by position between both the blood and fibroblast clones. **A** Individual 1: left hip (D1-L-H), Right hip (D1-R-H1 and D1-R-H2), left forearm (D1-L-F), and right forearm (D1-R-F1 and D1-R-F2). **B** Indivdual 2: left hip (D2-L-H), Right hip (D2-R-H), left forearm (D2-L-F), and right forearm (D2-R-F).

**Supplementary Figure 2:**
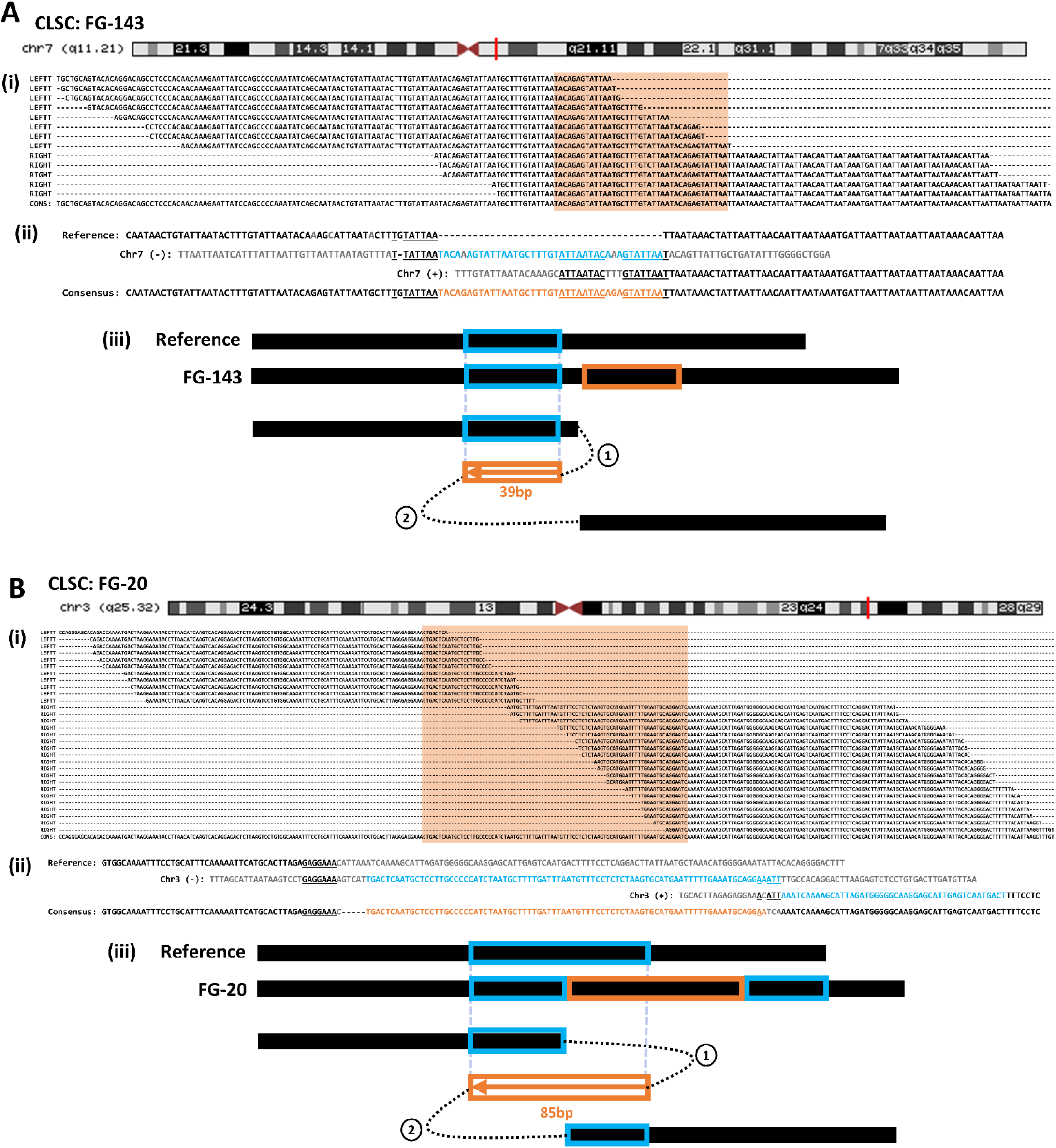
Examples of Classic MMBIR events found in germline of Fibroblast samples. **A** Classic MMBIR (CLSC) FG-143 found on q11.21 band of Chromosome 7 of individual D1. **(i)** Read cluster output generated by MMBSearch for FG-143. Cluster contains a 39bp insertion (orange box). **(ii)** Alignment of event FG-143 consensus to reference template. The insertion is shown in orange text and the template in light blue. Microhomology used in each template switching event is underlined. **(iii)** Schematic illustrating template switching events that lead to the formation of FG-143. Circled numbers indicate the order of template switches. Boxed arrows indicate the direction of synthesis, numbers under boxes indicate the length of synthesis and colors correspond to those in **(ii)**. **B** CLSC event FG-20 found on q25.32 band of Chromosome 3 of individual D2. **(i)** Read cluster containing an 85bp insertion (orange box). **(ii)** Alignment of event FG-20 consensus to reference template. Insertion is shown in orange text with its template (light blue). Microhomology is underlined. **(iii)** Schematic illustrating template switching events that lead to the formation of FG-20. Circled numbers indicate the order of template switches. Boxed arrows indicate direction of synthesis, numbers under boxes indicate length of synthesis, and colors correspond to those in **(ii)**.

**Supplementary Figure 3:**
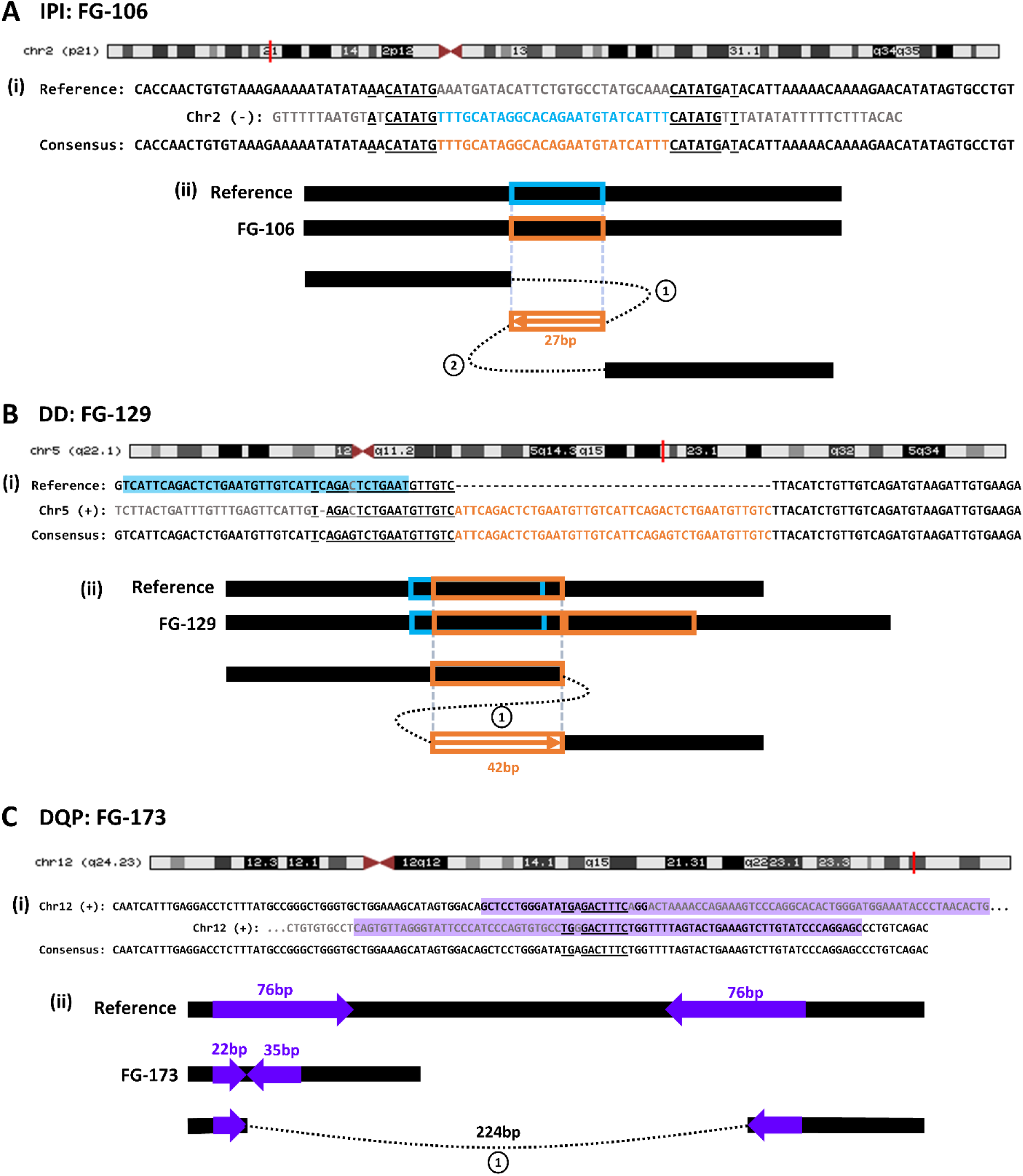
Examples of several classes of MMBIR events found in germline of Fibroblast samples. **A** Representative In-place Inversion (IPI) event FG-106 found on p21 band of Chromosome 2 of individual D1. **(i)** Alignment of event FG-106 consensus to reference template. Insertion (orange text) is 27bp and completely replaces the template (light blue text) end-to-end. Microhomology is underlined. **(ii)** Schematic illustrating one possible configuration of template switches. Circled number indicates the order of template switches. Boxed arrow indicates direction of synthesis, numbers under boxes indicate length of synthesis, and colors correspond to those in **(i)**. **B** Representative Direct Duplication (DD) event FG-129 found on q22.1 band of Chromosome 5 of individual D1. **(i)** Alignment of event FG106 consensus to best reference template. Insertion (orange text) is 42bp and best aligns to a forward template (also in orange text). The inverted template found by MMBSearch is highlighted in light blue. Microhomology is underlined. **(ii)** Schematic illustrating one possible configuration of template switches. Circled number indicates the single template switch. Boxed arrow indicates direction of synthesis, numbers under boxes indicate length of synthesis, and colors correspond to those in **(i)**, including the light blue box showing the location of the inverted template. **C** Representative Deletion at a Quasi-palindrome (DQP) event FG-173 found on q24.23 band of Chromosome 12 of individual D1. (i) Alignment of event FG-173 consensus to reference deletion junction locations. Palindromic sequences are highlighted in purple. Microhomology is underlined. **(ii)** Schematic illustrating the template switching event that resulted in a 224bp deletion between two 76bp inverted repeats (purple arrows). After the deletion, 22bp of the left repeat, and 35bp of the right repeat remained. Circled number indicates the single template switch. Colors correspond to those in **(i)**.

**Supplementary Figure 4:**
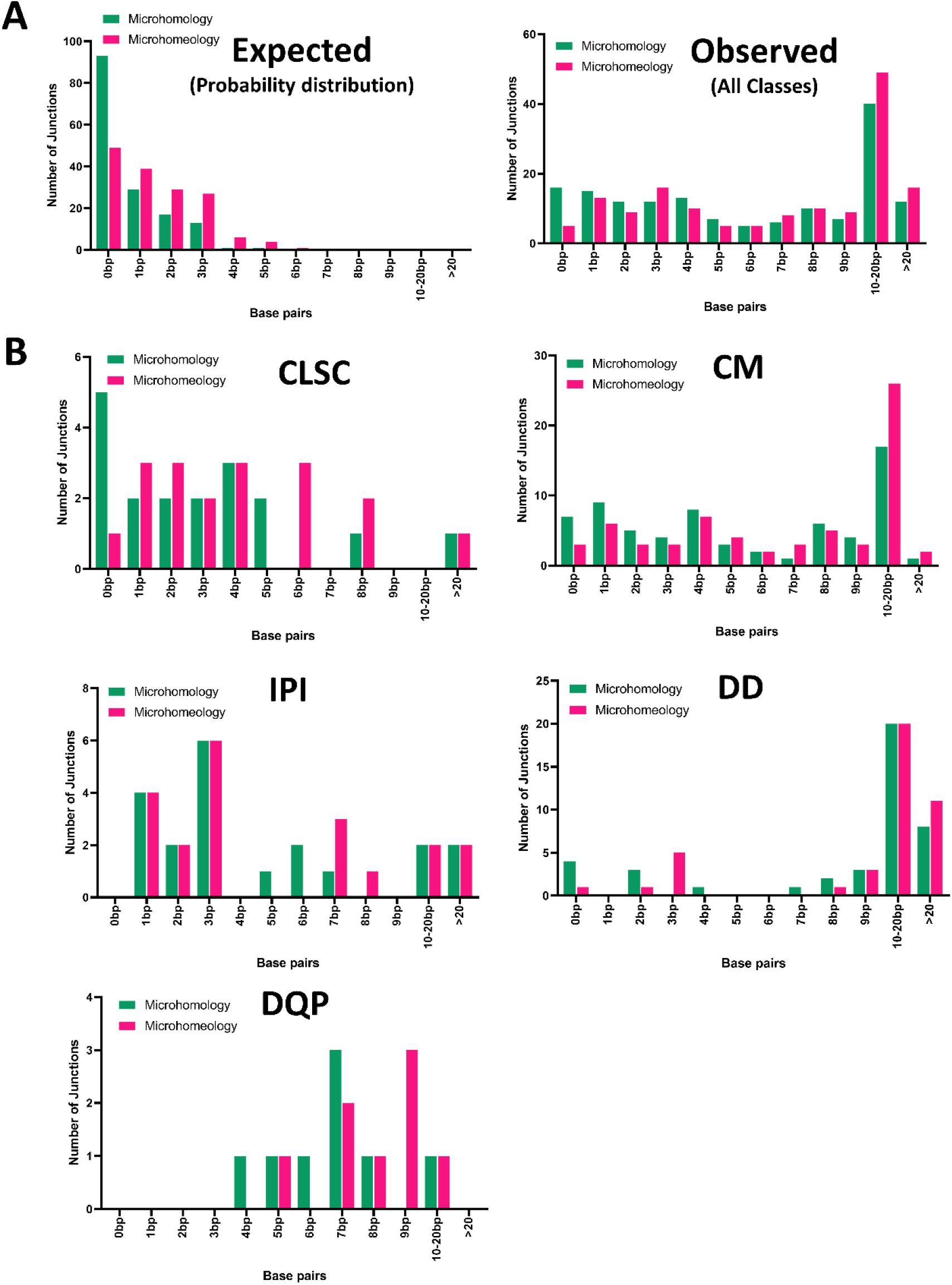
Microhomology distributions for all MMBIR classes found in germline of Fibroblast clones of individual D1. **A** Microhomology (Green bars) and Microhomeology (Pink bars) distributions for simulated MMBIR events (Expected) and all classes of MMBIR events found in germline of Fibroblast clone SRR4047717 (Observed). The “Expected” microhomology and microhomeology probability distribution is based on sampling of artificial randomly inserted MMBIR events on Chromosome 17 (data set used in Figure 1D) and multiplying by the total 155. D1 MMBIR junctions analyzed in “Observed”. **B** Distributions of microhomology and microhomeology for different classes of MMBIR events (CLSC, CM, IPI, DD, and DQP) that make up all classes shown in **A**(Observed) from individual D1 (See Supplementary Data S6).

**Supplementary Figure 5:**
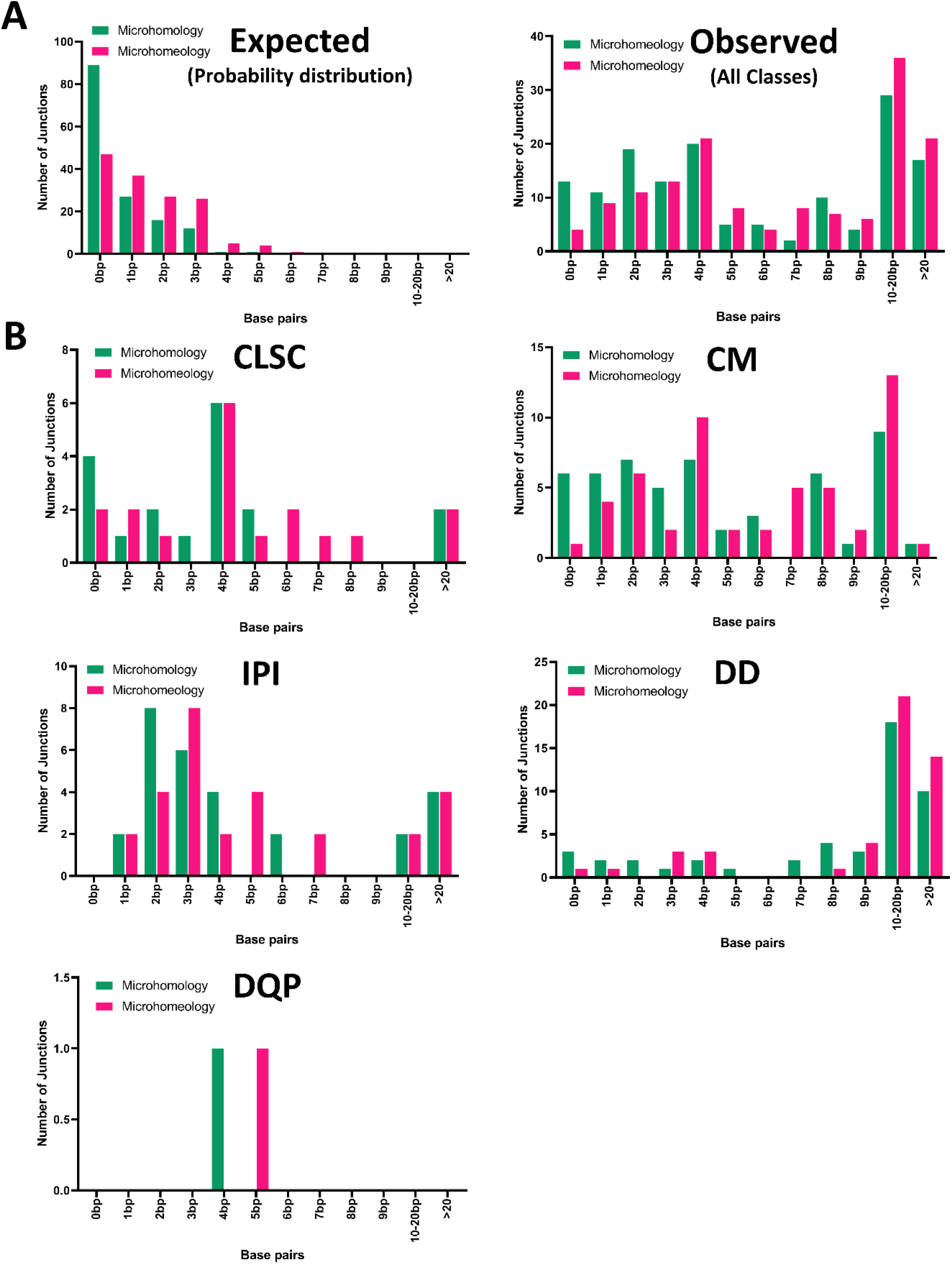
Microhomology distributions for all MMBIR classes found in germline of Fibroblast clones of individual D2. **A** Microhomology (Green bars) and Microhomeology (Pink bars) distributions for simulated MMBIR events (Expected) and all classes of MMBIR events found in germline of Fibroblast clone SRR4047718 (Observed). The “Expected” microhomology and microhomeology probability distribution is based on sampling of artificial randomly inserted MMBIR events on Chromosome 17 (data set used in Figure 1D) and multiplying by the total 148. D1 MMBIR junctions analyzed in “Observed”. **B** Distributions of microhomology and microhomeology for different classes of MMBIR events (CLSC, CM, IPI, DD, and DQP) that make up all classes shown in **A**(Observed) from individual D1 (See Supplementary Data S4).

**Supplementary Figure 6:**
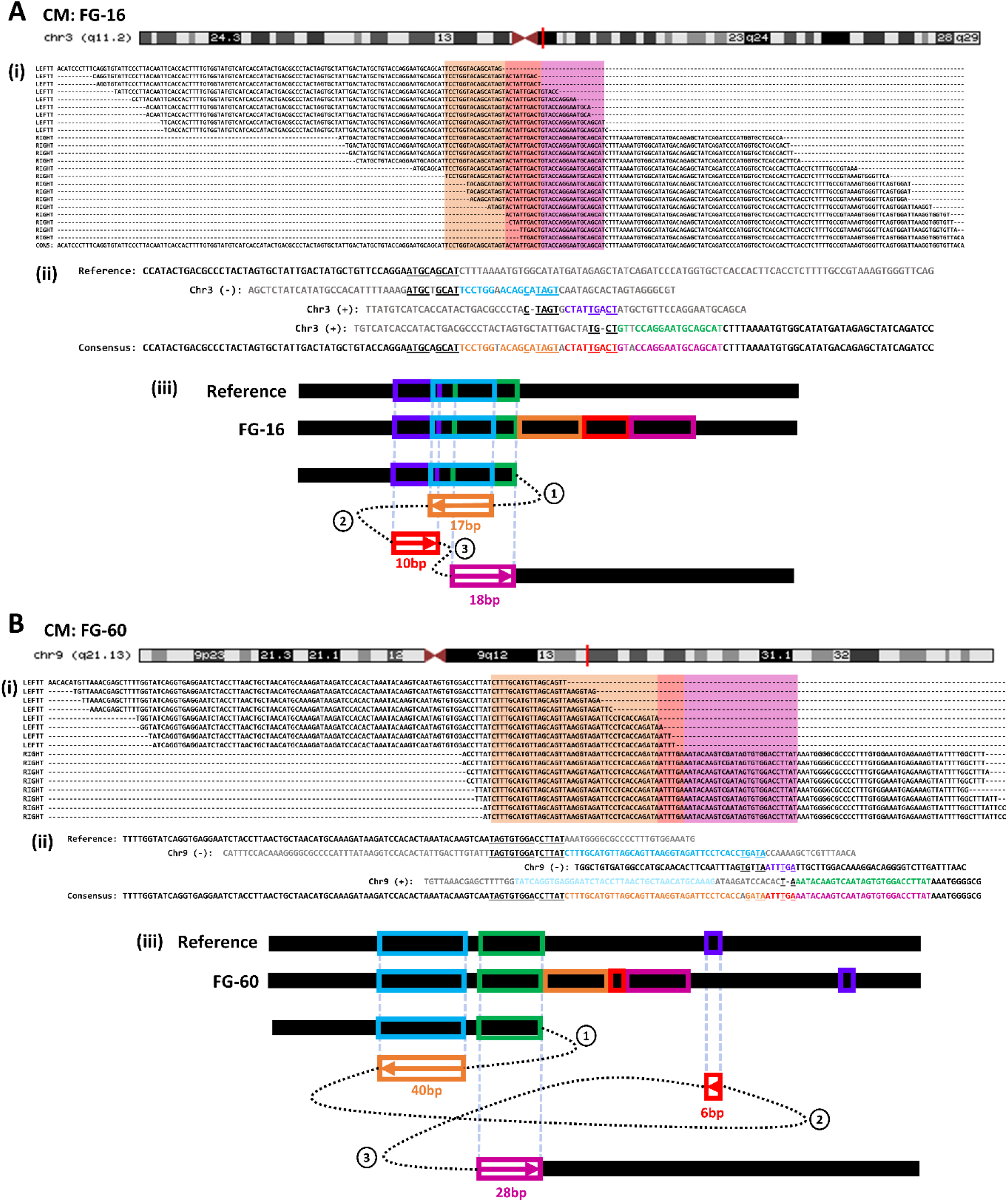
Examples of Complex MMBIR events found in germline of Fibroblast samples. **A** Complex MMBIR (CM) event FG-16 found on q11.2 band of Chromosome 3 of individual D2. **(i)** Read cluster output created by MMBSearch for FG-16. Read cluster contains a 45bp insertion which consists of 3 parts, each copied from different templates (distinguished by 3 colored boxes). **(ii)** Alignment of event FG-16 consensus to all refence templates. Insertion consists of 3 parts (orange, red, and pink text) copied from all corresponding templates (light blue, purple, and green respectively). Microhomology used in each template switching event is underlined. **(iii)** Schematic illustrating template switching events that lead to the formation of FG-16. Circled numbers indicate the order of template switches. Boxed arrows indicate direction of synthesis, numbers under boxes indicate length of synthesis, and colors correspond to those in **(ii)**. **B** CM event FG-60 found on q21.13 band of Chromosome 9 of individual D2. **(i)** Read cluster output generated by MMBSearch for FG-60. Read cluster contains a 73bp insertion which consists of 3 parts, each copied from different templates (distinguished by 3 colored boxes). **(ii)** Alignment of event FG-60 consensus to all reference templates. Insertion consists of 3 parts (orange, red, and pink text) copied from all corresponding templates (light blue, purple, and green respectively). Microhomology used in each template switching event is underlined. **(iii)** Schematic illustrating template switching events that lead to the formation of FG-60. Circled numbers indicate the order of template switches. Boxed arrows indicate direction of synthesis, numbers under boxes indicate length of synthesis, and colors correspond to those in **(ii)**.

**Supplementary Figure 7:**
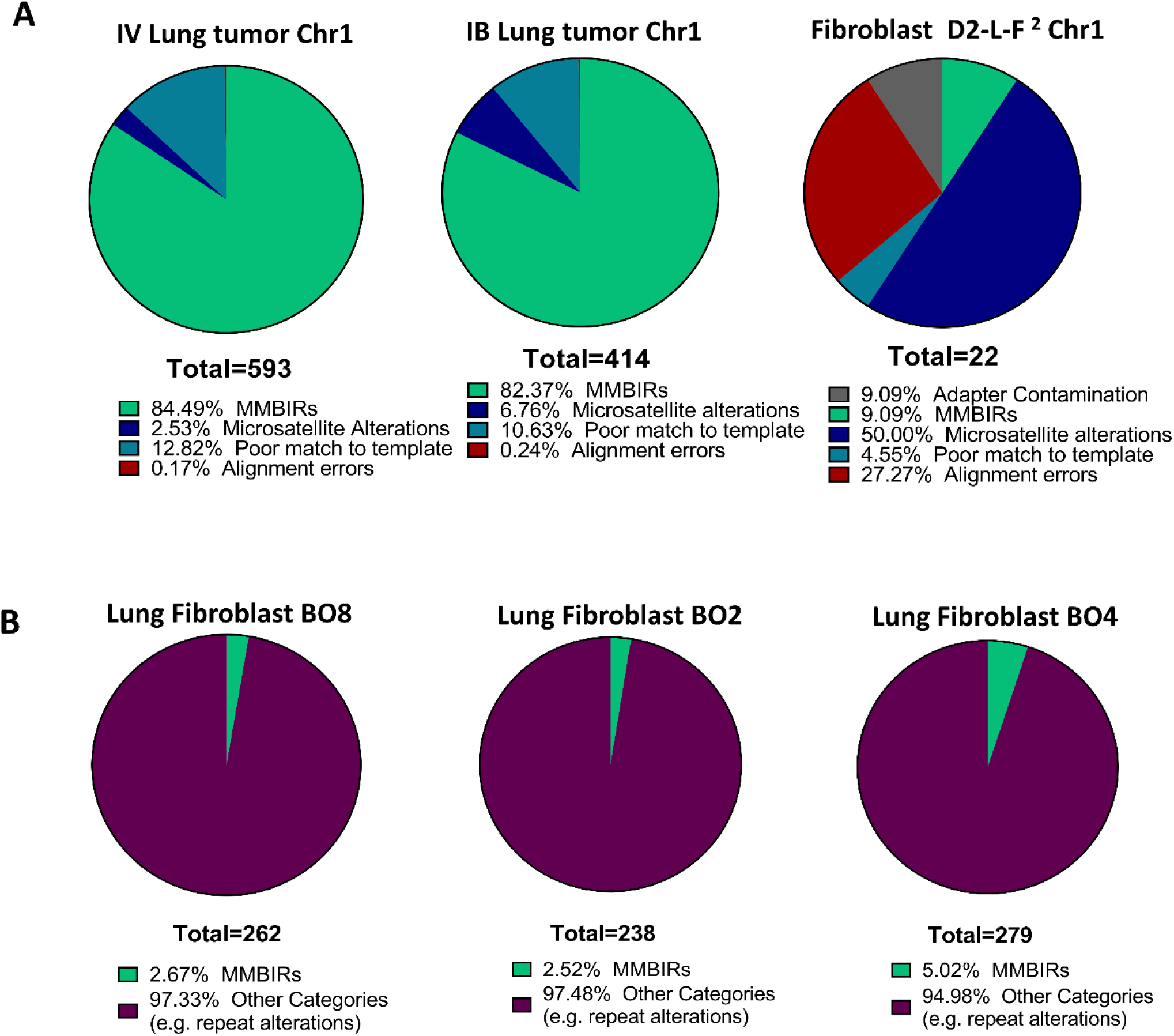
Comparison of MMBSearch calls from Chromosome 1 of lung tumors, and fibroblast and cultured lung fibroblast controls. **A** Analysis of Chromosome 1 MMBSearch calls (from clusters with ≥3 reads, insertions that are ≥10bp in length, and ≥80% identity between template and insertion) for IV Lung tumor (BO13-UIBB-1654), IB Lung tumor (SRR556475) obtained from dbGaP, and Fibroblast D2-L-F (SRR4047705) from ^2^. Germline MMBSearch calls were excluded for these sets. “MMBIRs” are those that resemble the Classic MMBIR pattern (CLSC). **B** Analysis of Chromosome 1 MMBSearch calls (parameters identical to **A**) for 3 Lung fibroblast clones BO8, BO2, and BO4. All MMBSearch calls were analyzed without exclusion of common calls. “MMBIRs” are those that resemble the Classic MMBIR pattern (CLSC).

**Supplementary Figure 8:**
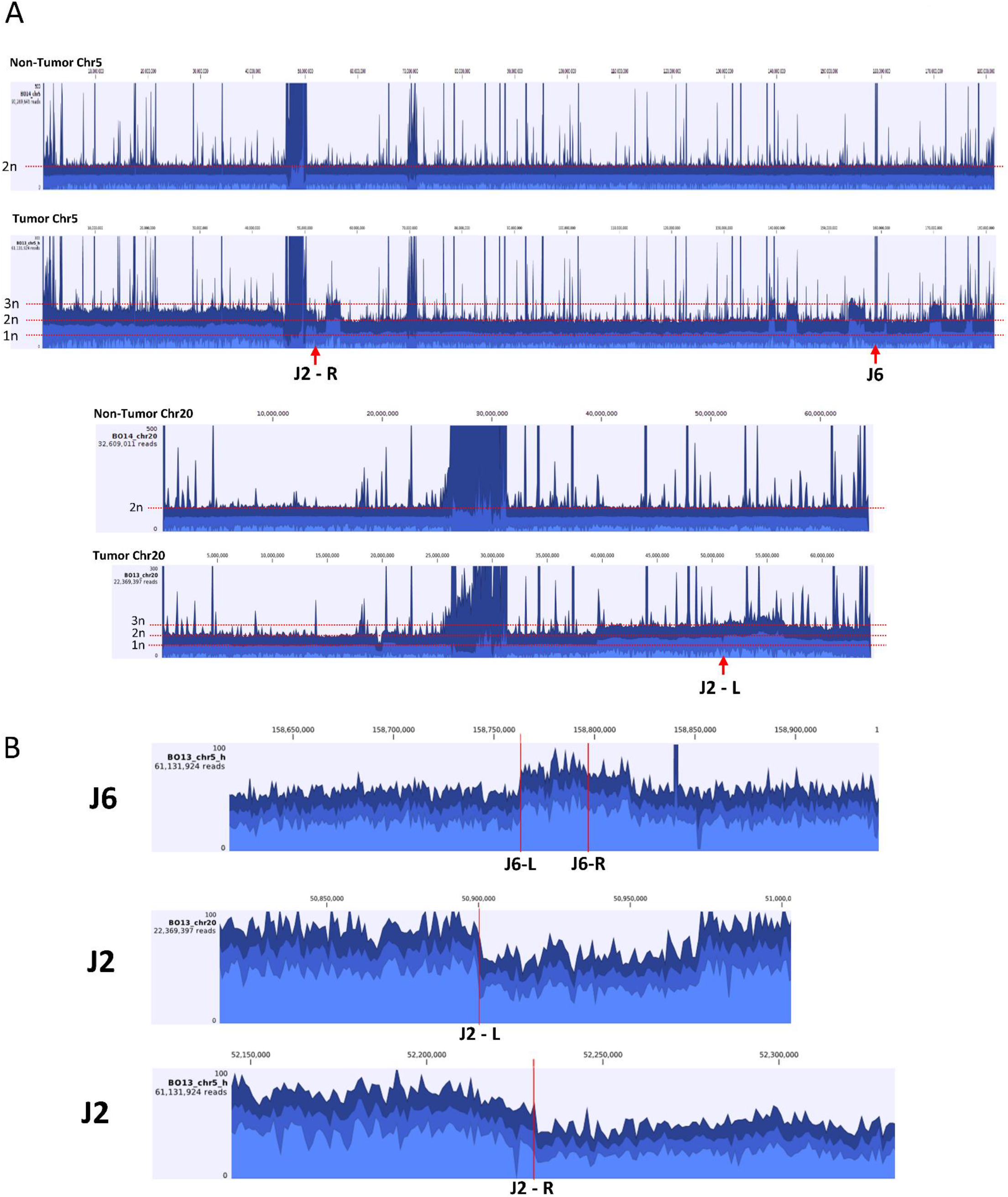
Read aggregation maps show copy number increases on Chromosomes 5 and 20 in IB Lung tumor sample. **A** Read aggregation tracks for entire Chromosomes 5 and 20 of tumor and non-tumor samples from IB Lung tumor. Locations of MMBIR junctions J2 (R: right side, L: left side) and J6 are indicated with red arrowheads. Read aggregation levels for different ploidies are indicated with red dotted lines. **B** Zoomed-in read aggregation tracks for the locations of J6 and J2 (R: right side, L: left side) in tumor sample, indicated as red lines.

## Supplementary Data

**Supplementary data S1.** MMBSearch analysis of fibroblasts of individual D1 from ^2^.

**Supplementary data S2.** MMBSearch analysis of fibroblasts of individual D2 from ^2^.

**Supplementary data S3.** MMBSearch output for Fibroblast-specific MMBIR call with analysis.

**Supplementary data S4.** Analysis of germline events and microhomology in individual D2 from SRR4047718 (D2-R-F) from ^2^.

**Supplementary data S5.** Maps of all germline MMBIR events found in sample SRR4047718 (D2-R-F) from ^2^.

**Supplemental data S6.** Analysis of germline events and microhomology in individual D1 from SRR4047717 (D1-L-F1) from ^2^.

**Supplementary data S7.** Maps of all germline MMBIR events found in sample SRR4047717 (D1-L-F1) from ^2^

**Supplementary data S8.** Summary of sequencing and MMBSearch Calls for tumor (BO13-UIBB-1654) and matched non-tumor (BO14-UIBB-1654) samples from stage IB lung adenocarcinoma patient.

**Supplementary data S9.** MMBSearch output for All chromosome 1 tumor-specific results for sample BO13-UIBB-1654.

**Supplementary data S10.** Analysis of all tumor-specific MMBsearch calls on chromosome 1 from sample BO13-UIBB-1654.

**Supplementary data S11.** Clonal and sub-clonal tumor-specific MMBIR and SV junctions from sample BO13-UIBB-1654.

**Supplementary data S12.** Primers for confirming Tumor-specific (BO13-UIBB-1654) junctions of selected MMBIRs and SVs by PCR and Sanger sequencing.

**Supplementary data S13.** Maps of breakpoint junctions listed in supplementary data S11.

**Supplementary data S14.** Summary of sequencing and MMBSearch Calls for tumor (SRR556475) and matched non-tumor (SRR551334) samples from stage IV lung adenocarcinoma patient from dbGaP.

**Supplementary data S15.** Analysis of all tumor-specific MMBsearch calls on chromosome 1 from sample SRR556475.

**Supplementary data S16.** MMBSearch output for all chromosome 1 tumor-specific results for sample SRR556475.

**Supplementary data S17.** Tumor-specific complex MMBIR junctions from sample SRR556475.

**Supplementary data S18.** List of all TCGA samples analyzed by MMBSearch.

**Supplementary data S19.** Summary of analysis of sample-specific MMBSearch calls from TCGA cancer genomes.

## Supplementary text

### Testing MMBSearch sensitivity with artificially generated read sets

To test the sensitivity of the MMBSearch in finding the MMBIR pattern, we created a set of synthetic genomes that each contained 993 MMBIR insertions 20-50 bp in size on human Chr 17 (See Methods for details). From these synthetic genomes, we generated 125bp paired-end artificial Illumina sequencing reads with an average insert size of 500bp and amounting to 30x average coverage across chromosome 17 using the NGS read simulator ART (Huang, Li, Myers, & Marth, 2012). After analyzing these read sets with the MMBSearch, Recall (sensitivity) and false positives were calculated for each length and type of insertion by comparing the MMBIR insertions called by the program to the list of insertion sequences within a 10% tolerance (e.g. 5bp for a 50bp insertion). Recall was highest for longer insertions that did not replace sequence (92-97% for insertions without replacement, and 65-77% with replacement) (Figure 1C, D), which is likely because the MMBSearch leverages reads that differ greatly from the reference and relies on local alignment to accurately differentiate between the insertion and its flanking reference-matching sequence, making more disruptive insertions easiest to identify and more likely to have their insertions called in full. Likewise, shorter insertions and those that replace sequence are more prone to partial calls. The number of false positives (see Methods) detected by the program was between 8 and 34 for various insertion lengths, representing 1-3% of calls made by MMBSearch. These false positives are caused solely by misalignment of half-reads to some repetitive regions of chromosome 17.

### Description of sub-clonal complex MMBIR events in lung cancer from dbGaP

By surveying Chromosomes 1-7, we found 41 sub-clonal events (defined as being present in fewer reads than 30% of the average read coverage for the sample) that, in addition to an insertion copied from a nearby template, contained sequence copied from a separate template (Supplementary Data S17), making them similar in structure to the events shown in Figure 6. For these complex events, 14 of the secondary templates were identified using BLAT as distant from the first template, resulting in chromosomal rearrangements and typically fusions with other chromosomes (Supplementary Data S17). For example, one MMBIR event that mediated a fusion between Chr 7 and Chr X (LT-1) (Figure 6E) contained 4bp of microhomology at its first junction, copied 19bp from Chr7 and using 9bp of microhomology, proceeded to Chromosome X. These junctions could be minimally extended as microhomeology with only a single base mismatch or gap. The second MMBIR event (LT-2) that mediated a fusion between Chromosome 7 and Chromosome 6 contained 5bp (extended to 7bp of microhomeology), copied 26bp from Chromosome 7, and used 5bp of microhomology to continue to Chromosome 6 (Figure 6F). Interestingly, additional matching bases were found further behind 3 of the 4 junctions in these examples (Figure 6E, F – yellow boxes).

### Analysis of fibroblast and cultured GM1604 cells with MMBSearch set to low clustering threshold

We reanalyzed one fibroblast sample from (Saini et al., 2016), with identical parameters to those used to call MMBIR events in the lung tumor sample that we sequenced (BO13-UIBB-1654). The analysis found only 22 MMBSearch calls unique to Chr1 in the fibroblast (Supplemental figure 7A) and only 3 of the calls appeared to match the MMBIR signature when manually inspected, while the remaining were either microsatellites or alignment errors. It is important to note that the fibroblast sample from (Saini et al., 2016)was prepared using a comparable library preparation kit to that which was used to prepare our lung tumor sample, except that it included a short PCR amplification step prior to sequencing. Our lung tumor sample by contrast was prepared using a PCR-free library prep kit to avoid accumulating palindromic artifacts introduced during end-blunting which have been shown to be preferentially amplified during library construction that utilizes PCR in highly fragmented samples (Star et al., 2014). Together, the MMBIR events called sub-clonally in the fibroblast sample were extremely rare.

As a secondary control, we performed sequencing on cultured GM1604 human fetal lung fibroblast cells. We chose this cell line as a control because it is non-cancer and originates from the lung. We selected single cells from the cultures and grew clonal lineages prior to sequencing (See Methods for culturing details). These cultured cells were sequenced using the same PCR-free library preparation as the lung tumor sample, and the same sequencing method (see Methods for details). We analyzed the sequencing reads from the GM1604 clones with MMBSearch and found that on Chr1 there was a low level (2.5-5%) of calls that matched the MMBIR signature (Supplementary Figure 7B). Together we conclude that the number of MMBIR insertions observed in both fibroblasts and the GM1604 clones was extremely low as compared to what was observed in both our sequencing of IB lung tumor, and IV lung tumor sequencing obtained from dbGaP (Supplementary Figure 7A), supporting our idea that the high quantity of MMBIR events found in the lung tumor samples is related to the pathology of the tumor and not a technical artifact.

